# Intracellular calcium stores mediate a synapse-specific form of metaplasticity at hippocampal dendritic spines

**DOI:** 10.1101/460568

**Authors:** Gaurang Mahajan, Suhita Nadkarni

## Abstract

Long-term plasticity mediated by NMDA receptors supports input-specific, Hebbian forms of learning at excitatory CA3-CA1 connections in the hippocampus. An additional layer of stabilizing mechanisms that act globally as well as locally over multiple time scales may be in place to ensure that plasticity occurs in a constrained manner. Here, we investigate the potential role of calcium (Ca^2+^) stores associated with the endoplasmic reticulum (ER) in the local regulation of plasticity dynamics at individual CA1 synapses. Our study is spurred by (1) the curious observation that ER is sparsely distributed in dendritic spines, but over-represented in large spines that are likely to have undergone activity-dependent strengthening, and (2) evidence suggesting that ER motility within synapses can be rapid, and accompany activity-regulated spine remodeling. Based on a physiologically realistic computational model for ER-bearing CA1 spines, we characterize the contribution of IP_3_-sensitive Ca^2+^ stores to spine Ca^2+^ dynamics during activity patterns mimicking the induction of long-term potentiation (LTP) and depression (LTD). Our results suggest graded modulation of the NMDA receptor-dependent plasticity profile by ER, which selectively enhances LTD induction. We propose that spine ER can locally tune Ca^2+^-based plasticity on an as-needed basis, providing a braking mechanism to mitigate runaway strengthening at potentiated synapses. Our model suggests that the presence of ER in the CA1 spine may promote re-use of synapses with saturated strengths.

## Introduction

Hebbian synaptic plasticity involves activity-driven changes in synaptic strengths^1^. This form of plasticity is inherently unstable, as a small change in synaptic strength can promote further change in the same direction, and this positive reinforcement can drive synaptic efficacies to either saturate or shrink to a minimu^2-4^. It has long been recognized that Hebbian rules need to be supplemented with additional stabilizing mechanisms to curb runaway plasticity and support stable yet flexible neural circuit^2, 5, 6^. The issue of stability is usually addressed within the theoretical Bienenstock-Cooper-Munro (BCM) framework^7^, which posits an adaptive threshold for long-term potentiation induction that varies as a function of the history of prior activity of the postsynaptic neuron, concurrently affecting all its afferent synapses. There is, however, limited understanding of the principles governing biophysical implementations of such a rule^8-13^. This instantiates the more general question as to what physiological mechanisms exist, at the cellular and synaptic levels, that could actively regulate the balance of plasticity and stability through appropriate adjustment of the rules of activity-induced synaptic alterations, thereby shaping the long-term dynamics of modifiable synapses^14^.

Synaptic plasticity, a potential neural substrate for learning and memory storage, is particularly well studied in the hippocampal formatio^15, 16^, and much about its molecular underpinnings has been learned from investigations at the excitatory CA3 to CA1 Schaffer collateral (SC) synapse, an integral component of the neural circuitry supporting the encoding of spatial location (place fields^)17-20^. These synapses are capable of undergoing bidirectional modification (both long-term potentiation (LTP) and long-term depression (LTD)) which relies primarily on the activation of postsynaptic N-methyl-D-aspartate/glutamate receptors (NMDAR) and ensuing calcium (Ca^2+^) entry into the dendritic spin^21-24^. Given the biophysical requirements of both membrane depolarization and glutamate binding for their activation^25^, NMDAR are naturally poised to mediate input-specific, Hebbian forms of synaptic learnin^26, 27^. However, an account of Ca^2+^ signaling based solely around transmembrane receptors is incomplete, as another source of Ca^2+^ may be available to CA1 spines via Ca^2+^ release from the endoplasmic reticulum (ER)^28^. ER, which is present throughout the dendritic arbor of hippocampal neurons, has a heterogeneous distribution in individual spines^29^. Curiously, its occurrence is skewed towards the larger spine heads: nearly 80% of the larger mushroom-shaped spines show presence of ER, but overall, ER extends into only about 20% of all spines in adult dendrites, as seen in EM serial reconstruction of rat CA1 pyramidal cell dendrites^30^.

Here, we consider a synapse-specific form of metaplasticity that may be introduced by Ca^2+^ stores associated with the ER in these large dendritic spines. Our investigation is motivated by the observation that larger spines that have ER are associated with stronger synapses and have most likely been potentiate^31-34^. Further, recent imaging studies on cultured hippocampal neurons from mice reveal a dynamic picture of the ER distribution in spines^35^; ER can undergo rapid growth in individual spine heads on a timescale of minutes, which was shown to be regulated by NMDAR activation^36^, and accompanies spine enlargemen^37, 38^. This local remodeling of the ER in spines correlated with changes in spine morphology suggests an adaptive function for ER specifically at potentiated synapses, providing an interesting context to our investigation.

Several lines of previous experimental work indicate an involvement of the intracellular ER store in neuronal Ca^2+^ regulation^39^, with possible implications for activity-regulated plasticity processe^40-42^. This, however, still leaves open important questions regarding the role of stores in microdomain signaling in the context of its uneven distribution in CA1 spines, and, given that small spines without ER are capable of undergoing potentiation, what additional functionality it introduces in the context of plasticity at individual synapses. Ca^2+^ release from ER is particularly associated with Ca^2+^ signaling underlying synaptic long-term depression^15^. Experimental studies on hippocampal/cortical LTD have implicated a requirement for group I metabotropic glutamate receptor (mGluR) signaling (particularly involving the mGluR5 subtype) in the induction of depression by prolonged low frequency synaptic stimulatio^43, 44^, and this was shown to depend on release of Ca^2+^ from inositol 1,4,5-triphosphate (IP_3_)-sensitive intracellular store^45-47^. These earlier findings were based on recordings of synaptic population responses; thus, pointed questions on the synapse specificity of mGluR signaling and Ca^2+^ store contribution could not be directly addressed. Two subsequent imaging-based studies on plasticity at individual synapses provide more clarity in this regard. Glutamate uncaging at individual CA3-CA1 synapses in rat hippocampal slices^32^ was reported to evoke restricted Ca^2+^ release from ER in individual spine heads via IP_3_ signaling, and mGluR/store-dependent depression induced by prolonged low frequency uncaging stimulation was found to be localized to synapses associated with such ER-bearing spines. Results from a more recent similar study of spine structural plasticity^48^ show that, at synapses associated with larger spines, contribution of mGluR/IP_3_-mediated store Ca^2+^ release in these spines is necessary for the induction of synaptic weakening with low frequency input trains. Together, the above findings suggest that, at least at a subset of synapses that are likely to have undergone experience-dependent potentiation, IP_3_-mediated Ca^2+^ release from spine ER may make a particularly significant contribution under weak stimulation, augmenting the NMDAR-mediated Ca^2+^ influx to facilitate synaptic depression.

Computational models of hippocampal synapses typically attempt to account for experimental findings on LTP/LTD in terms of the postsynaptic Ca^2+^ elevation mediated by varying levels of NMDAR activatio^49-53^. These models provide a useful description of a canonical, or generic, synapse. Given that presence of spine ER may be restricted to a small proportion of synapses, the contribution of ER is likely to be ‘washed-out’ and difficult to disambiguate based on coarse-grained population readouts. Clear description of Ca^2+^ signaling by ER and its distinctive effects on plasticity in CA1 spines does not exist despite suggestive evidence that synaptic plasticity can be modulated by the contribution from internal stores. To address this conspicuous gap in our understanding, we have undertaken a detailed characterization of the engagement of the ER Ca^2+^ store at an active CA3-CA1 synapse. Our modeling study of an ER-bearing CA1 dendritic spine head integrates a detailed kinetic description of mGluR-regulated Ca^2+^ release from ER with a realistic model for NMDAR Ca^2+^ signaling. Our analysis highlights the synapse-level differences in Ca^2+^ signaling and metaplasticity arising from presence of ER, and provides a novel perspective on the functional role of ER, as an intracellular Ca^2+^ store, in the postsynaptic context.

## Results

### A calibrated kinetic model for mGluR-IP_3_ receptor signaling recapitulates salient features of experimental data on Ca^2+^ release from ER in dendritic spines

We consider an average CA1 pyramidal neuron dendritic spine head containing ER (an ER+ spine), described by a single-compartment point model in the well-mixed approximation (Methods). The time course of spine Ca^2+^ evoked by synaptic activation is shaped by the coupled electrical and Ca^2+^ dynamics at the spine head, which involves contributions from various biophysical components, and passive electrical coupling with a dendritic compartment (Figs. 1A-1C; Tables S1-S4). The spine ER store, modeled as an intracellular Ca^2+^ pool (fixed luminal concentration of 250 μM), contributes through IP_3_ receptor-gated Ca^2+^ release (IP_3_ and Ca^2+^-induced Ca^2+^ release, or ICCR) and the uptake of cytosolic Ca^2+^ by SERCA pumps present on the ER membrane.

**Figure 1.**
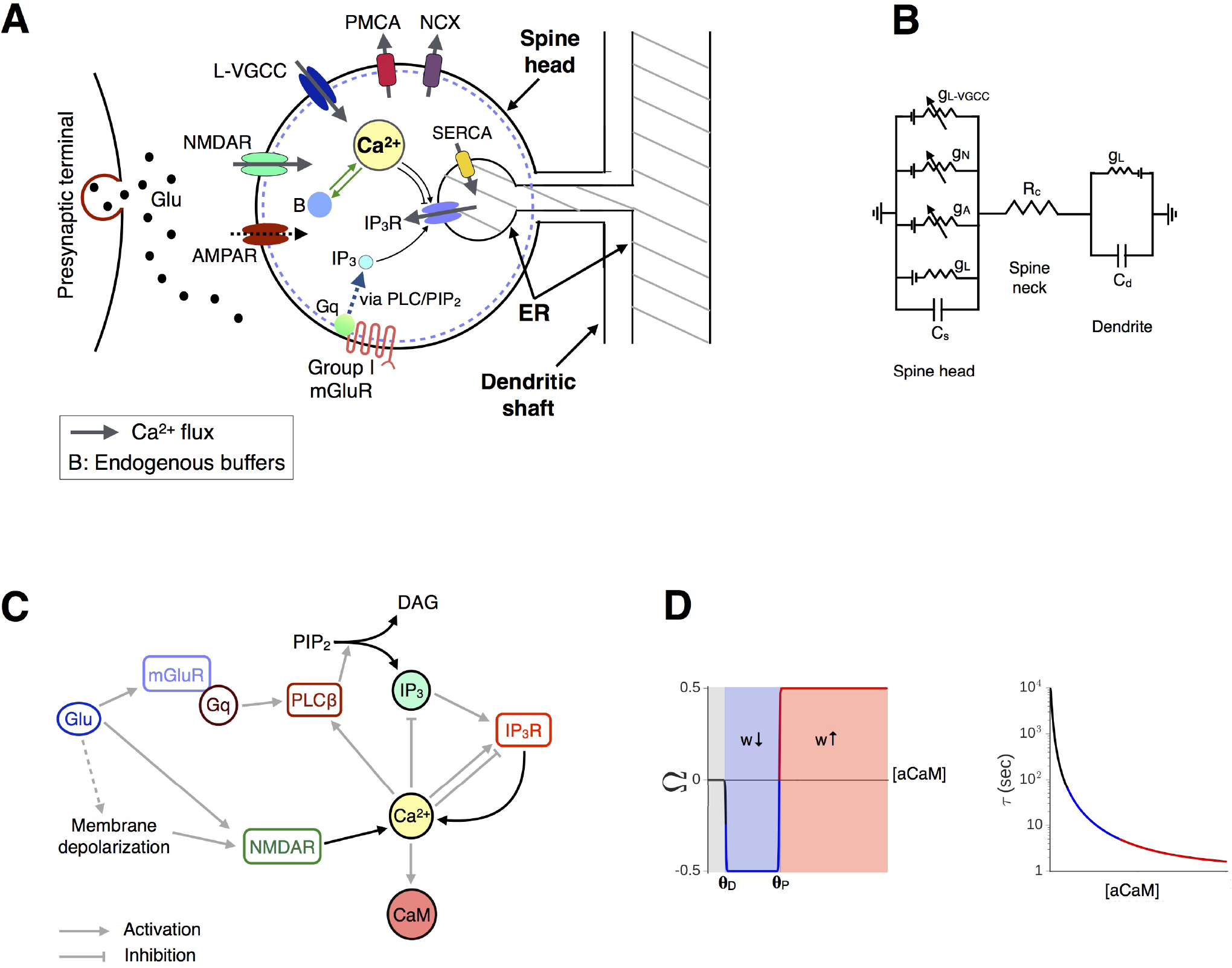
Modeling activity-driven Ca^2+^ dynamics and plasticity at a dendritic spine with ER. **A**, Schematic representation of an ER+ hippocampal CA1 dendritic spine head outlining the various model components included in this study. Glutamate release into the synaptic cleft activates postsynaptic AMPA and NMDA receptors along with group I metabotropic receptors (mGluR), driving Ca^2+^ increase in the spine due to NMDAR-gated entry from outside the cell and Ca^2+^ release from the IP_3_-sensitive ER store. **B**, Equivalent circuit diagram representing the spine head compartment which is resistively coupled to a (passive) dendritic compartment via the neck resistance R_*c*_. **C**, Summary of the biochemical cascade involved in IP_3_-and Ca^2+^-mediated Ca^2+^ release (ICCR) from the ER store, which contributes to (and is potentially influenced by) spine Ca^2+^ rise mediated by NMDAR. Ca^2+^ interacts with calmodulin (CaM), which regulates the activity of downstream effectors governing spine plasticity. **D**, The functions Ω and τ encapsulating the Ca^2+^/calmodulin (CaM)-based synaptic plasticity model used in the present study (Methods). Ω is parametrized by the thresholds *θ_D_* and *θ_P_*, which control the windows for depression (blue) and potentiation (red) of the weight variable *w*.

In order to have a model of mGluR-IP_3_ signaling and ICCR that is appropriate for describing hippocampal synapses, we refer to Ca^2+^ imaging data from a previous experimental study of long-term depression at individual excitatory CA3-CA1 connections^32^. Flash photolysis of caged glutamate was reported to evoke mGluR-and IP_3_-dependent store Ca^2+^ release which was specific to ER+ spines, and trailed the initial NMDAR-mediated spine Ca^2+^ transient (trial-averaged delay of ∼470 ms). We adopted a detailed kinetic scheme^54^ to describe the sequence of biochemical events linking the initial binding of glutamate to postsynaptic Gq-coupled receptors (expressed on the extrajunctional membrane) to the eventual synthesis of IP_3_ and DAG via PIP2 hydrolysis mediated by activated PLCβ (Methods). We tuned the parameters regulating IP_3_ production rate in this model (Table S3) to reproduce the empirical estimate for the timing of the second Ca^2+^ peak relative to the arrival of glutamate (we chose parameters such that for a synapse with *N_R_* = 30 IP_3_ receptors and Δ*Ca_EPSP_* ~ 0.2*μ*M, the second Ca^2+^ peak occurs with a delay of t ~ 480 ms). Figs. 2A and 2B show the simulated Ca^2+^ response to application of a single pulse of glutamate at *t* = 0 in our model spine head with ER (colored curves), which is compared with the reference ER-spine (black curve). The latency as well as magnitude of Ca^2+^ release from ER are dependent on the number of IP_3_ receptors present. Although Ca^2+^ at low levels (≲ 0.3 *μ*M) is a coagonist for IP_3_R activation^55^, release of ER store Ca^2+^ in our calibrated model does not require the initial NMDAR-mediated Ca^2+^ transient (Fig. S1), which, again, is consistent with experimental findings^32^.

**Figure 2.**
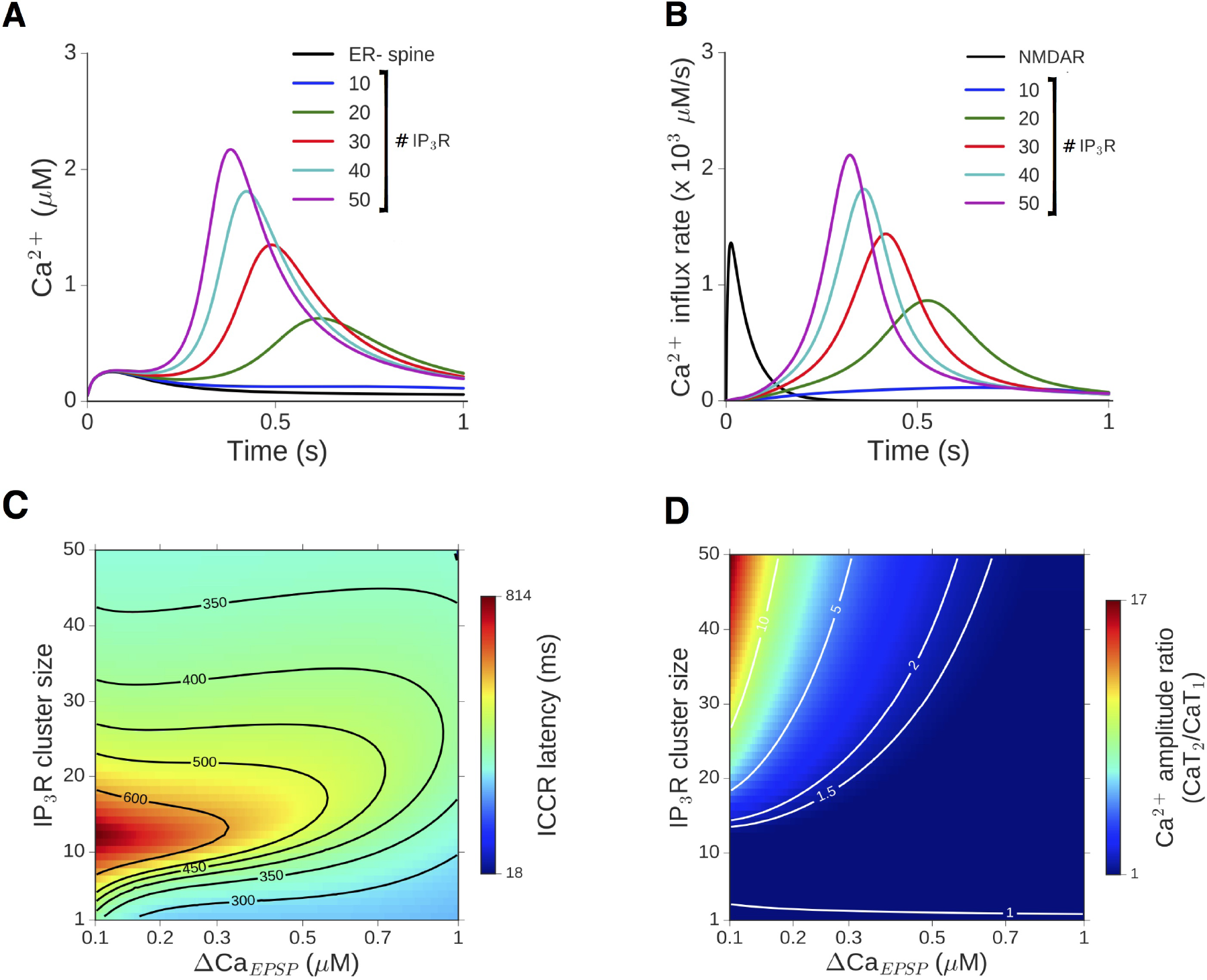
Glutamate evokes delayed release of Ca^2+^ from dendritic spine ER via mGluR signaling and IP_3_R activation. **A**, Response of the reference ER-spine (black) to a single glutamate input is compared with the Ca^2+^ time course in ER+ spines with 10-50 IP_3_ receptors (colored curves). The peak of the NMDAR-mediated Ca^2+^ rise is Δ*Ca_EPSP_* = 0.2 *μ*M. **B**, The underlying Ca^2+^ flux through NMDAR channels (black) and the delayed flux through IP_3_R channels (colored curves). **C**, Dependence of the delay in ICCR in the ER+ spine on the number of IP_3_ receptors and amplitude of the initial NMDAR-evoked Ca^2+^ (Δ*Ca_EPSP_*). The horizontal axis is linear in the NMDAR conductance parameter *g*_N_, but parameterized by Ca^2+^ for ease of interpretation; contours correspond to various values of the latency. **D**, Dependence of ratio of the second to the first Ca^2+^ peak (illustrated in A) on the number of IP_3_ receptors and Δ*Ca_EPSP_*. Contours shown for different values of the amplitude ratio.

The amplitude of the synaptically evoked Ca^2+^ (a direct readout for the NMDAR conductance) and the IP_3_ receptor cluster size directly determine the Ca^2+^ signal in the spine. Changes in ICCR profile in our calibrated model corresponding to changes in these crucial components is summarized as heat maps in Figs. 2 C & D. These represent the dependence of the delay of ICCR (in ms) and the amplitude of the second Ca^2+^ peak relative to the first, respectively, over a biologically realistic range of parameter values. These parameter ranges yield a fairly broad distribution of possible outcomes, and only for a small subset of parameter choices, the outcomes are consistent with the averaged experimental estimates^32^ (latencies of μ400-500 ms and a Ca^2+^ ratio CaT_2_/CaT_1_ of 1-5). Notably, the delay in synaptically evoked release of store Ca^2+^ has a non-monotonic dependence on the IP_3_R cluster size: for a fixed Δ*Ca_EPSP_*, the delay first increases, and then decreases with increasing number of IP_3_R. This dependence may be understood by noting that the IP_3_R cluster size sets the strength of the Ca^2+^-dependent positive feedback loop driving IP_3_R activation in the presence of sufficient IP_3_. For higher IP_3_R numbers, this feedback can drive a self-sustained burst of Ca^2+^ release from ER. The larger the cluster size, the more swiftly ICCR increases, while also speeding up the suppression of the slower Ca^2+^-dependent inactivation variable h which eventually shuts off the IP_3_R channel flux. Thus, the temporal profile of the ICCR transient is expected to advance (shorter lag) with increasing IP_3_R number. At small cluster sizes, on the other hand, the flux through IP_3_R is inadequate to drive a burst of ICCR even while IP_3_ is present. In this regime, there is no discernible second Ca^2+^ peak (e.g., the blue curve corresponding to 10IP_3_R in Fig. 2A), and the kinetics of the IP_3_R open fraction (i.e., the ICCR peak location) in this case is primarily shaped by the decaying NMDAR Ca^2+^ transient. The relatively minor contribution of ICCR only adds a small delay to this decay of the total spine Ca^2+^. This additional delay is proportional to the IP_3_R number, N_R_, thus accounting for the shift of the peak location of IP_3_R flux to longer latencies with increasing N_R_ in the regime of small IP_3_R numbers.

### mGluR-mediated Ca^2+^ release from ER can facilitate the induction of synaptic depression with weak stimulation

We next simulate the calcium response in an ER+ spine head during the induction of synaptic weakening by low frequency afferent stimulation. Experimental blockade of group I mGluR signaling or depletion of IP_3_-sensitive Ca^2+^ is shown to compromise synaptically-induced LTD at SC synapses onto CA1 pyramidal cell^43, 45, 47, 48^. Fig. 3A shows an example of the Ca^2+^ time course evoked by a 1 Hz train of regularly spaced glutamate pulses. Binding of glutamate to postsynaptic AMPA receptors (AMPAR) produces a small depolarization at the spine head (~few mV), resulting in weak NMDAR activation and modest Ca^2+^ entry. Due to little overlap of successive Ca^2+^ events at low input rates, there is no build-up of Ca^2+^ concentration in the spine over time. Fig. 3A compares the Ca^2+^ signal in the ER-control spine (black curve) with the responses in the ER+ spine (colored curves correspond to different numbers of IP_3_R). mGluR-mediated Ca^2+^ release from spine ER contributes to the common pool of Ca^2+^ in the spine head and augments the NMDAR-mediated Ca^2+^ signal.

**Figure 3.**
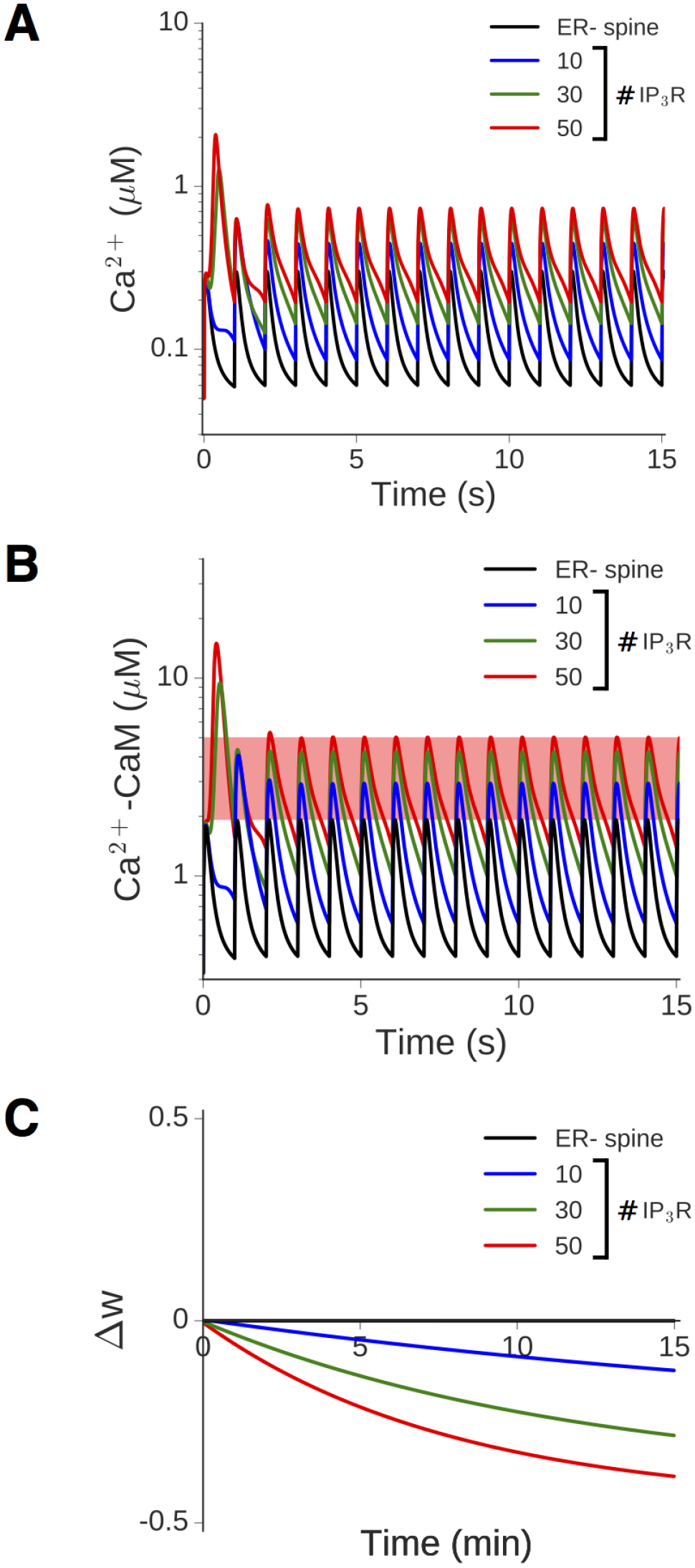
Ca^2+^ release from ER enhances the spine Ca^2+^ signal at low input rates and facilitates LTD induction. **A**, Time course of Ca^2+^ evoked by repeated synaptic input at 1 Hz in the model ER-spine (black) and an equivalent ER+ spine with 10-50 IP_3_ receptors. Δ*Ca_EPSP_* = 0.2 μM in the ER-spine. **B**, The corresponding time dependence of Ca^2+^-bound calmodulin (CaM) in the ER-/ER+ spine head. Total CaM concentration = 50 μM. **C**, Change in the synaptic weight variable, *w*, driven by the activated CaM during a 1 Hz input train applied for 15 minutes (900 spikes). The plasticity threshold *θ_D_* has been set to 2 μM, such that no LTD is induced in the absence of ER contribution, and LTD induction is facilitated with increasing flux through IP_3_R.

In order to map the Ca^2+^ time course to plasticity, we also follow the activation of calmodulin (CaM), which is included in our model and makes a significant contribution to the Ca^2+^ buffering capacity in the spine (Fig. 3B). Ca^2+^-bound CaM is known to regulate a number of downstream signaling molecules that collectively determine changes in synaptic strength associated with early LTP/LTD; thus, it provides a suitable choice of input to the dynamical model governing the synaptic weight *w* (Methods and Fig. 1D). The parameter θ_*D*_ in Eq.10 decides the Ca^2+^/CaM threshold for LTD induction. For the example in Fig. 3B, a range of choices is possible (indicated by the red band running parallel to the time axis) such that LTD can be induced at the ER+ spine, whereas the synaptic strength associated with the ER-spine remains unaffected. This is illustrated in Fig. 3C by a comparison between the time courses of *w*(*t*) in the ER-and ER+ spines over 900 SC inputs for the specific choice of *θ_D_* = 2 *μ*M. The enhanced Ca^2+^/CaM response in the ER+ spines (colored curves) drives a slow induction of synaptic depression (Δ*w*<0) with repeated stimulation, which is absent in the ER-spine (black curve). This particular instantiation of our model thus recapitulates experimental observations regarding the association of mGluR-mediated store Ca^2+^ release with LTD induction at low stimulation rate^47, 48^.

In order to address the dependence of the results on the model parameters, we have repeated our simulations across a range of synaptic input frequencies (0.1-5 Hz) and synaptically evoked spine Ca^2+^ amplitudes (Δ*Ca_EPSP_* = 0.1-1 *μ*M). Fig. 4A shows the dependence of the maximum steady-state amplitude of active CaM during persistent stimulation on the input frequency. The corresponding total change in the weight variable (Δ*w*) at the end of the stimulus train is shown in Fig. 4B (*f_D_* is fixed at 1 Hz). ICCR robustly enhances CaM activation to facilitate LTD induction over a range of low frequencies, although the contribution of ICCR is non-monotonic in the input rate. This dependence may be accounted for by noting that the opening of IP_3_R is regulated by a combination of two factors: the Ca^2+^-and IP_3_-dependent activation, and the level of inhibition (*h*). At low input rates (≲ 2 Hz), the IP_3_ level increases with the frequency of glutamate application. At the same time, the slowly changing *h* variable has less time to recover between successive inputs, the more frequently the inputs arrive; therefore, *h* decreases with increasing input frequency. The balance between these two competing factors (IP_3_-mediated activation and inactivation mediated by *h*) shapes the overall profile of the IP_3_R open probability, and hence the IP_3_R flux, as a function of the frequency of glutamate input.

**Figure 4.**
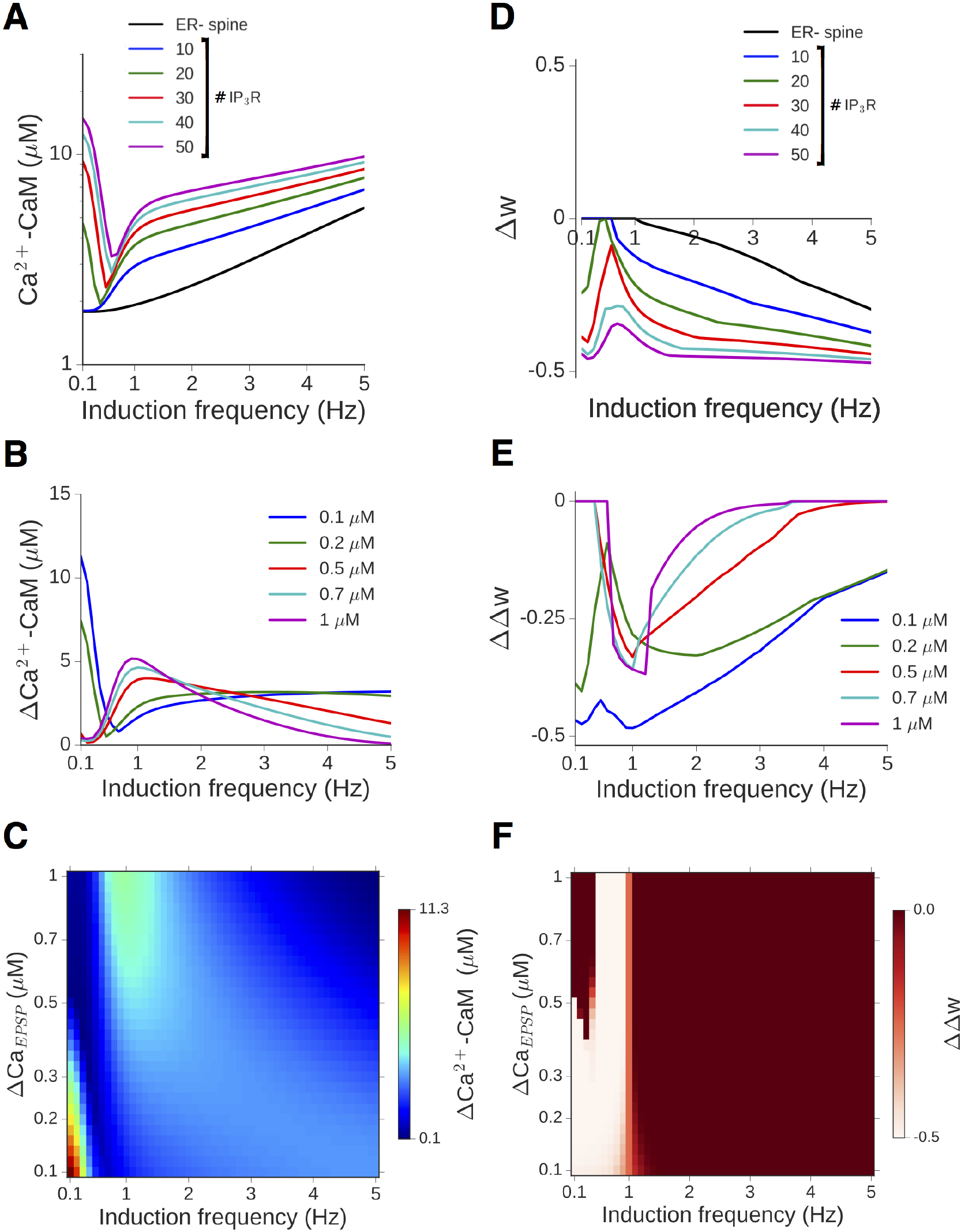
ICCR robustly enhances spine Ca^2+^/CaM response and facilitates LTD over a range of synaptic configurations and low frequency inputs. **A**, The amplitude of CaM response to synaptic inputs in the 0.1-5 Hz range is compared between the ER-spine (black) and equivalent ER+ spine with 10-50 IP_3_ receptors (colored curves). **B**, The differential CaM response profile in an ER+ spine with 30 IP_3_R relative to the ER-spine for different choices of DCaEPSP (i.e., NMDAR conductance). **C**, Dependence of the maximal CaM activation in the ER+ spine with 30 IP_3_R, relative to the ERspine, on the NMDAR conductance and input rate. **D**, Change in the weight variable *w* at the end of 900 spikes in the ER-(black) and ER+ spines (colored curves), corresponding to **A**. *θ_D_* is set to give no LTD in the ER-spine for frequencies ≤ 1 Hz. **E**, Differential LTD induction, ΔΔ*w*, in an ER+ spine with 30 IP_3_R relative to an ER-spine for different choices of Δ*Ca_EPSP_*. **F**, Dependence of the excess weight change (ER+ spine with 30 IP_3_R v/s ER-spine) on the NMDAR conductance and the input rate. Lighter colors indicate enhanced LTD response in the presence of ICCR.

Figs. 4C and 4D compare the responses in ER-/ER+ spines over a range of low frequencies for different choices of the NMDAR conductance parameter (Δ*Ca*_EPSP_), with a fixed number of IP_3_R (*N_R_* = 30). The profiles of the *excess* CaM activation (Fig. 4C) and differential plasticity outcome (Fig. 4C) in the presence of spine ER depend on the amplitude of synaptically evoked Ca^2+^. However, in general, ICCR contribution enhances LTD induction over a range of low-frequency inputs.

Elaborating on the above results, the heat maps in Figs. 4E and 4F summarize the dependence of the maximal Ca^2+^/CaM response and plasticity output in the ER+ spine, relative to the ER-reference spine, on the input frequency and NMDAR Ca^2+^ conductance. The vertical axes in both figures are linear in the NMDAR conductance parameter *g_N_*; however, to aid interpretation, they have been parametrized in terms of the NMDAR Ca^2+^ amplitude instead. As before, the frequency threshold for LTD induction is set to *f_D_* = 1 Hz, and all results are for an IP_3_R cluster size of *N_R_* = 30. ICCR is found to robustly enhance NMDAR-driven LTD induction at lower frequencies (f ≲ 1 Hz). The underlying CaM activation in our model exhibits complex dependence on the input rate and Δ*Ca_EPSP_*(Figs. 4B and 4C): the excess CaM response in the ER+ spine decreases with increasing Δ*Ca_EPSP_* at very low input frequencies (f ≲ 0.5 Hz), but this trend reverses at higher frequencies (f ~ 1-2 Hz).

To see why this difference arises, we first consider the case of very low frequencies (f ≲ 0.5 Hz). Due to the delayed synthesis of IP_3_, the initial NMDAR Ca^2+^ elevation only contributes to ICCR by changing the slower inactivation variable *h* (and not through the activation variable *m*_1_), which decreases with increasing Δ*Ca_EPSP_*. The level of *h* determines the magnitude of the subsequent ICCR (i.e. the amplitude of the second Ca^2+^ peak), and as *h* gets smaller with increasing Δ*Ca_EPSP_*, so does the maximal IP_3_R Ca^2+^ flux. At higher frequencies (~1-2 Hz), on the other hand, there is insufficient time between one input and the next for IP_3_ to decay back to resting levels. As IP_3_ is now present at moderate levels when a glutamate input arrives, the NMDAR-mediated Δ*Ca_EPSP_* switches its role: now it directly controls the activation of IP_3_R (through the instantaneous activation variable *m*_1_), while the slowly-evolving *h*(*t*) only has a delayed effect on the IP_3_R open probability. Due to this switch in its role from inactivation to activation at higher input rates, an increase in *g_N_*(i.e., Δ*Ca_EPSP_*) is now associated with increased IP_3_R flux. This explains the increase of ICCR with the NMDAR conductance. In sum, our results demonstrate that store Ca^2+^ contribution regulated by mGluR-IP_3_ signaling at a realistic synapse can robustly augment the NMDAR-mediated Ca^2+^ response to weak synaptic stimulation, providing a basis to understand compromised hippocampal LTD associated with blocking of ER Ca^2+^ stores.

### The contribution of ICCR to spine Ca^2+^ dynamics is regulated by NMDAR activation and depends on the rate of synaptic stimulation

Our experimentally constrained model of mGluR-IP_3_ signaling indicates that spine ER can make a substantial contribution to the Ca^2+^ response evoked by weak synaptic inputs. Next, we examine the input frequency dependence of the ER contribution to spine Ca^2+^ dynamics during persistent synaptic activation. The degree and polarity of long-term changes in synaptic efficacy are known to be controlled by the rate of SC pathway stimulation^56^. Higher input rates are associated with stronger NMDAR activation and greater build-up of Ca^2+^ in the spine head, leading to a switch from LTD to LTP induction above some crossover frequency *f_P_*. We wish to address how this rate-dependent bidirectional plasticity profile is modulated by localized Ca^2+^ release from spine ER.

A qualitative sense of how the gating of IP_3_R is regulated by the level of synaptic activation may be obtained from its dose-response profile, i.e., the steady-state open probability (*P_open_*) of IP_3_R as a joint function of glutamate and spine Ca^2+^ concentration. Increasing the level of glutamate stimulation (e.g., in the context of rate-based plasticity) has a direct effect on IP_3_ synthesis via mGluR-PLCβ activation. At the same time, it also drives increased postsynaptic NMDAR activation. The resulting elevation of spine Ca^2+^ level can affect both the production (via the mGluR pathway) and degradation (via the Ca^2+^ dependence of IP_3_K activity) of IP_3_, besides directly controlling *P_open_* through the *m*_1_ and h variables (Methods). The broad strokes of this nonlinear dependence on glutamate and Ca^2+^ can be captured by the asymptotic steady-state response of the IP_3_R over a range of (constant) glutamate and Ca^2+^ levels. Fig. 5 represents the steady-state *P_open_* as a 3D surface plot. IP_3_R shows maximal activation for a Ca^2+^ concentration ∼0.3 *μ*M over a range of realistic glutamate levels, and is progressively inhibited with increasing Ca^2+^. Thus, in an ER+ spine, the contribution of ICCR is anticipated to diminish progressively with increasing NMDAR activation for a fixed glutamate signal (as in the case of STDP induction protocol). Given the weak dependence of IP_3_ on the glutamate level as suggested by Fig. 5, we can also anticipate a similar graded contribution of ICCR with increasing NMDAR-mediated Ca^2+^ elevation in the context of rate-dependent plasticity. The interplay between NMDAR-mediated Ca^2+^ entry, mGluR signaling, and IP_3_R gating suggested by the above picture is characterized in detail in our synaptic model with realistic parameter settings.

**Figure 5.**
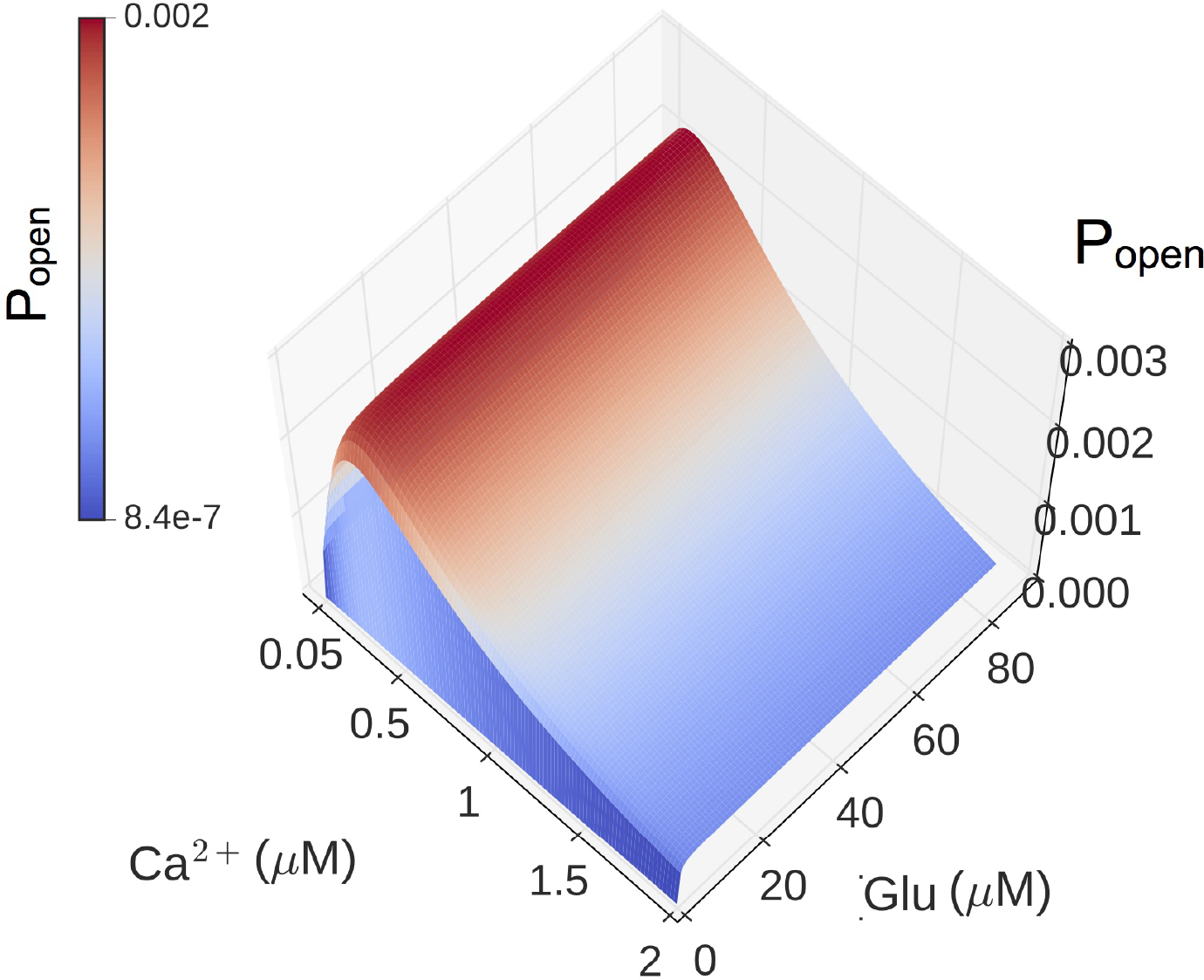
IP_3_ receptor activation is determined by glutamate and cytosolic Ca^2+^ levels. The steady state open probability of an IP_3_ receptor (*P_open_*) as a function of (constant) glutamate and Ca^2+^ concentrations.

We simulate a standard protocol consisting of repeated presynaptic stimulation (900 spikes) of the SC-CA1 pathway at different rates (0.1-20 Hz in steps of 0.1 Hz) (Methods). Fig. 6A compares the frequency-response profile of spine Ca^2+^ elevation (the maximum amplitude attained during the steady state) in the reference ER-spine (black) with the ER+ spine for different numbers of IP_3_R (colored curves). ICCR enhances the Ca^2+^ responses at lower frequencies, and its contribution steadily diminishes with increasing frequency above f ~ 5 Hz; this is also highlighted by the profile of the excess Ca^2+^ amplitude in the ER+ spine relative to the ER-spine, shown in Fig. 6B. Similar dependence on the input rate is also observed for the activation of CaM in the ER-and ER+ spines (Figs. 6C and 6D). The dip in the differential Ca^2+^ signal below zero at higher frequencies (Fig. 6B) is due to SERCA pump activity in the ER+ (but not the ER-) spine, which contributes to the extrusion of cytosolic Ca^2+^ and leads to a net lowering of the Ca^2+^ levels in the presence of ER. The action of SERCA Ca^2+^ efflux is revealed only at higher frequencies when ICCR in the spine is suppressed by the Ca^2+^-dependent inactivation of IP_3_R. The inverse dependence of ER Ca^2+^ contribution on the level of NMDAR activation is also highlighted by a direct comparison between the NMDAR and IP_3_R Ca^2+^ current profiles in the ER+ spine, which are shown in Fig. 6E.

**Figure 6.**
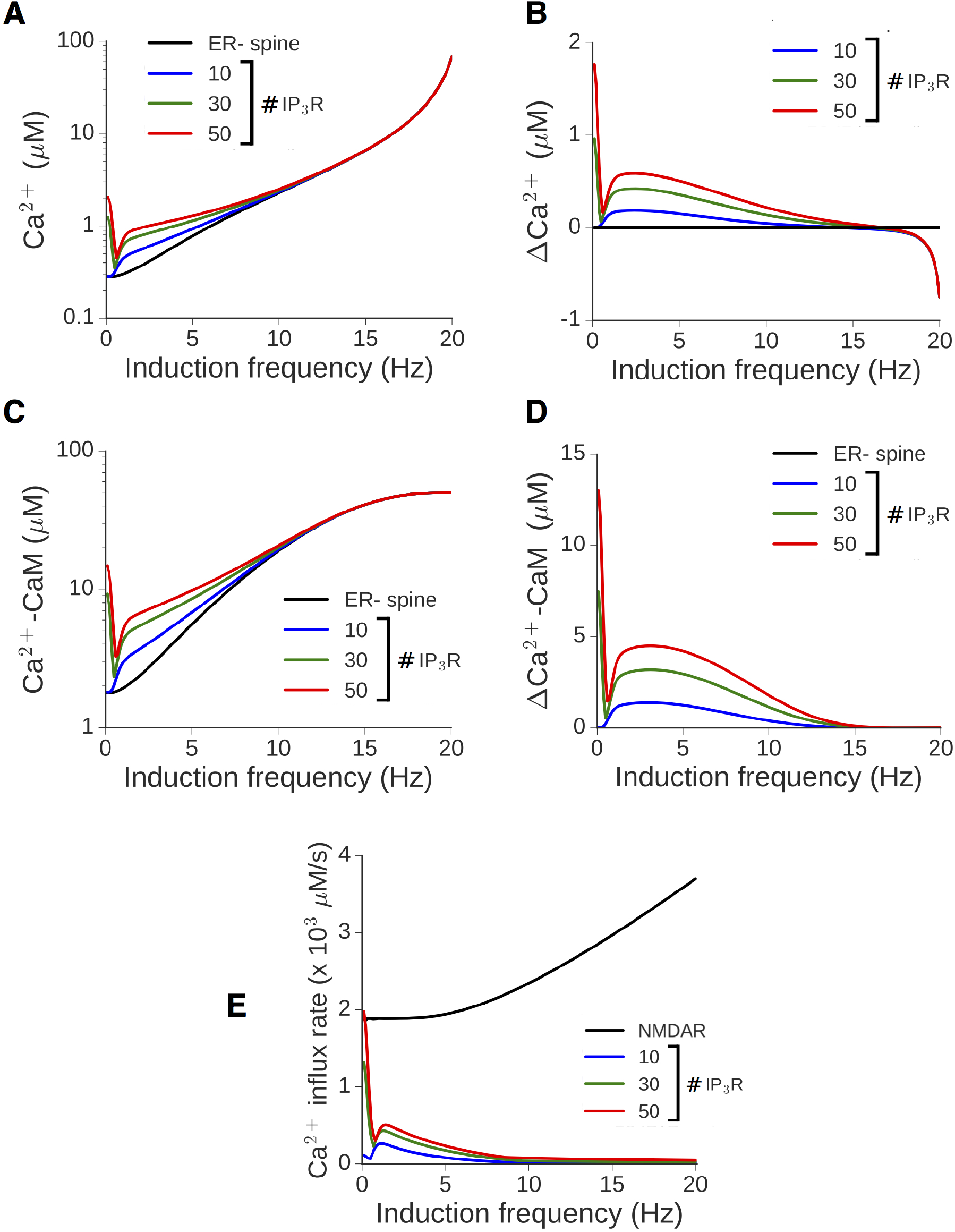
Augmented Ca^2+^/CaM response in a spine head in the presence of ER depends on the synaptic input rate and is suppressed at higher frequencies. **A**, Maximum Ca^2+^ level attained during persistent stimulation at different frequencies in the reference ER-spine (black) and equivalent ER+ spine with different numbers of IP_3_R (colored curves). **B**, Non-monotonic dependence of the differential Ca^2+^ responses in the ER+ spine on the input rate. **C, D**, The corresponding results for CaM activation as a function of the input rate. **E**, The maximum calcium influx rate through NMDA receptors (black) and different numbers of IP_3_ receptors (colored curves) in an ER+ spine plotted against the input rate. (All results for the model synapse with peak NMDAR-mediated Ca^2+^ response Δ*Ca_EPSP_* = 0.2 *μ*M.)

Spine Ca^2+^ elevation drives a change in the synaptic weight variable w (Eq. 10). This is illustrated for the ER-(control) spine in Figs. 7A and 7B, which show the temporal profile of Ca^2+^/CaM (Fig. 7A) and the corresponding evolution of the w variable (Fig. 7B) over 900 spikes at two different input rates, 5 and 17 Hz, associated with LTD and LTP, respectively. (The plasticity thresholds have been adjusted to have *f_D_* = 1 Hz and *f_P_* = 15 Hz at the ER-spine.) The differential effect of ER is displayed separately for the 5 Hz (Figs. 7C and 7D) and 17 Hz (Figs. 7E and 7F) examples. At 5 Hz, ICCR makes an appreciable contribution to the spine Ca^2+^ elevation, and thus to the rate and amplitude of the resulting synaptic changes, with a larger number of IP_3_R associated with stronger synaptic depression (Fig. 7D). In contrast, at 17 Hz, due to the strong suppression of ICCR by the NMDAR-driven persistent Ca^2+^ elevation, there is little difference in the response in the ER-and ER+ spines (Fig. 7E), resulting in nearly the same plasticity outcome (strong potentiation) at the end of the stimulation (Fig. 7F).

**Figure 7.**
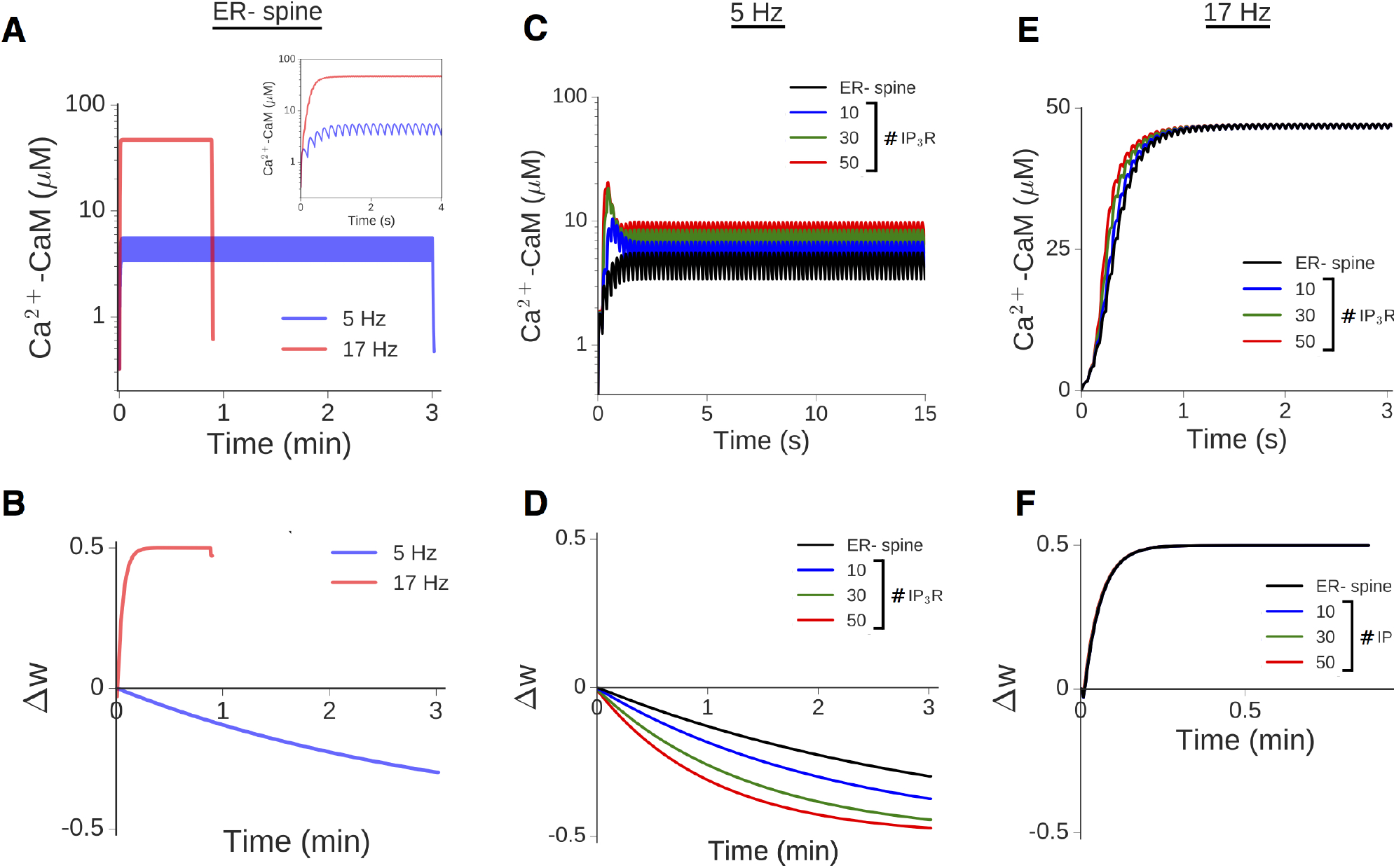
Rate-dependent Ca^2+^ elevation drives bidirectional plasticity at ER-/ER+ spines. **A,B**, Illustration of the CaM response (A) and plasticity induction (B) at the reference ER-spine for 900 presynaptic spikes applied at two different rates, 5 (blue) and 17 (red) Hz. Inset in A shows a magnified view of the first 4 seconds of the Ca^2+^/CaM time course. Plasticity parameters have been adjusted to yield LTD and LTP thresholds *f_D_* = 1 Hz and *f_P_* = 15 Hz, respectively. C,D, Enhancement of NMDAR-mediated CaM activation (C) and LTD induction (D) at 5 Hz stimulation due to ER Ca^2+^ contribution for different IP_3_R cluster sizes. **E,F**, Time course of CaM activation (E) and potentiation (F) at 17 Hz stimulation in the ER-spine (black) and ER+ spine for different IP_3_R cluster sizes (colored curves). Due to the suppression of ICCR at higher frequencies, presence of ER makes little difference in this case. In C and E, only the initial phase of the (much longer) time course is shown for clarity.

We have quantified the induced plasticity profile across the full range of synaptic input rates (0.1-20 Hz), which is shown in Fig. 8A. The total synaptic weight change (Δw) at the end of the stimulus train is plotted as a function of the input rate *f* for the reference ER-spine (black) and for the ER+ spine with different numbers of IP_3_R (colored curves). Consistent with our expectation from Fig. 5, we find that ICCR enhances plasticity at lower input frequencies, leading to a broadening of the effective LTD window. Due to the Ca^2+^-dependent suppression of ICCR at higher frequencies, the profiles of Δw for synapses associated with the ER-and ER+ spines are near-identical above f ~ 10 Hz.

**Figure 8.**
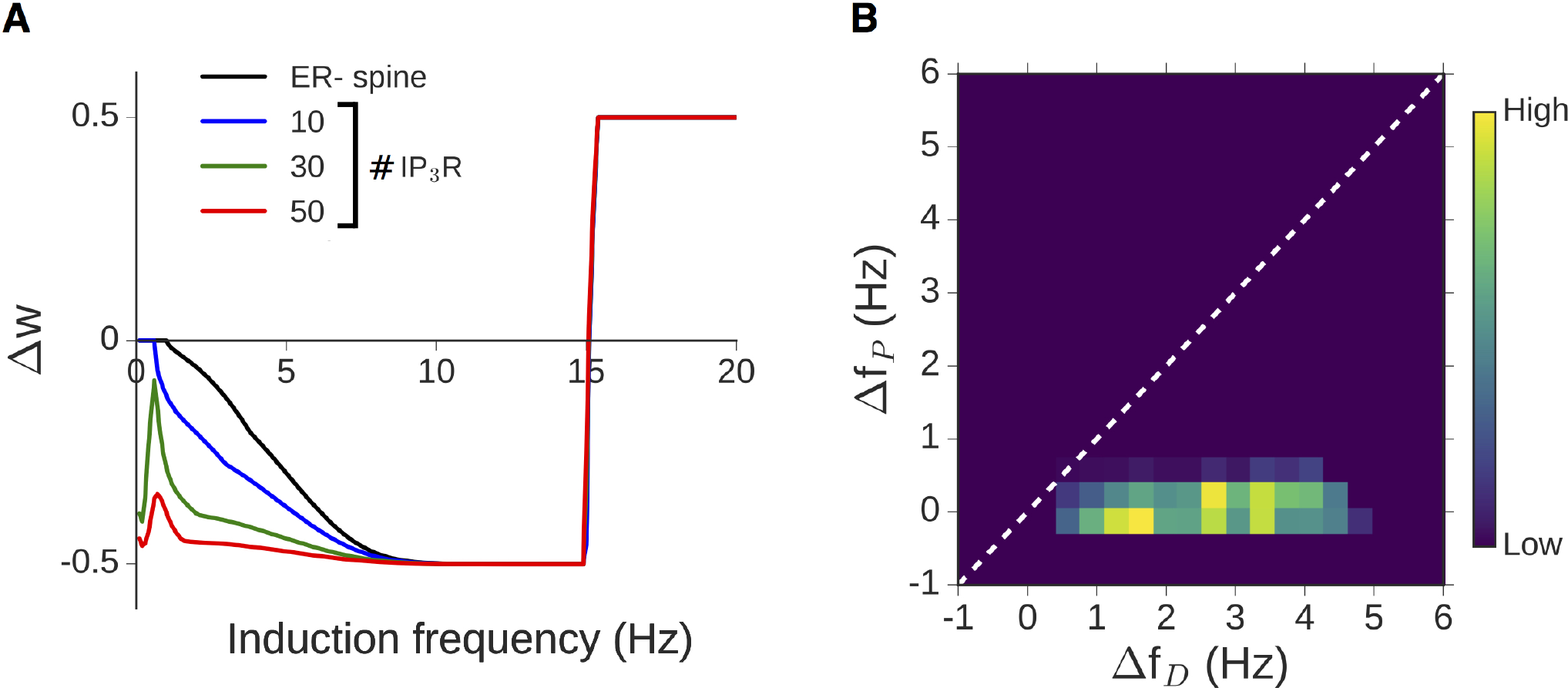
Graded contribution of ICCR to spine Ca^2+^ modulates the rate-dependent plasticity profile. A, Dependence of the total weight change (Δ*w*) induced by prolonged stimulation (900 spikes) on the synaptic input rate at the ER-spine (black) and corresponding ER+ spine for different IP_3_R numbers (colored curves). LTD/LTP thresholds are set to *f_D_* = 1 Hz and *f_P_* = 15 Hz. Plasticity is enhanced in the ER+ spine at low frequencies, with diminished modulation at higher (LTP-inducing) rates. B, Graded contribution of ICCR to the modulation of plasticity is quantified in terms of the relative shifts in the LTD/LTP thresholds in the ER+ spine (relative to the ER-control spine). The heat map is a distribution of Δ*f_D_*/Δ*f_P_* values obtained by random sampling (5000 times) of the plasticity thresholds *f_D_* and *f_P_* (over 1-6 Hz and 10-20 Hz, respectively), and IP_3_R number (10-50). (All results for a synapse with NMDAR-mediated Δ*Ca_EPSP_* = 0.2 μM.)

The differential modulation of LTD and LTP suggested by the above results for our model synapse (Δ*Ca_EPSP_* = 0.2 μM) is characterized in terms of the relative broadening of the depression and potentiation windows. A simple way to capture the overall modulation of the plasticity curve is by estimating changes in the threshold frequency for LTD induction (Δ*f_D_*) and LTP induction (Δ*f_P_*) with the inclusion of ER, relative to the ER-spine. The plasticity thresholds for the ER-spine (*f_D_* and *f_P_*) were repeatedly sampled at random from 1-6 Hz for *f_D_* and 10-20 Hz for *f_P_* to account for experimental uncertainties in these estimates, as well as assess the variability in the model output. The resulting distribution of (Δ*f_D_*, Δ*f_P_*) values (aggregate of 5000 runs over 10-50 IP_3_R) is visualized as a heat map in Fig. 8B, and on the whole, it suggests selective enhancement of LTD induction in the presence of ER. In summary, our analysis of the model spine head suggests a graded, frequency-dependent contribution of ER to spine Ca^2+^ signaling and plasticity, with steadily diminishing contribution of ICCR at higher input frequencies.

### Ca^2+^ release from the IP_3_-sensitive ER store selectively enhances synaptic depression during spike timing-dependent plasticity (STDP)

We next examine the involvement of ER in spine Ca^2+^ dynamics during trains of pre-and postsynaptic action potentials (APs). Experiments on hippocampal as well as cortical synapses suggest that the relative timing of pre-and postsynaptic spiking regulates the activation of NMDAR, and the resulting postsynaptic Ca^2+^ elevation directly controls the size and polarity of synaptic change^57-59^. In general, transmitter release preceding (trailing) the arrival of a somatically generated backpropagating action potential (BAP) at the postsynaptic spine is associated with synaptic strengthening (resp. weakening); however, the frequency of the paired stimulation also plays an important role in deciding the form of plasticity. In the context of hippocampal CA3-CA1 synapses, pairing of every presynaptic spike with 1 BAP at theta frequency (5 Hz) was found to induce only synaptic weakening, and LTP induction requires pairing glutamate release with AP bursts (2 BAPs) in the postsynaptic neuron^60^. We make use of this experimental data on STDP, specific to the CA3-CA1 synapse, to constrain our model for spine Ca^2+^ signaling, and examine the contribution of ICCR during this form of plasticity.

We have simulated the Ca^2+^ dynamics in our synaptic model during a sequence of pre-and postsynaptic spikes. Every glutamate input is separately paired with either 1 BAP (spike doublets) or 2 BAPs (spike triplets), and these paired stimuli are presented 100 times at a fixed rate of 5 Hz (Methods). Figures 9A and 9B illustrate the time course of Ca^2+^-CaM (Fig. 9A) and the corresponding change in the synaptic weight (Fig. 9B) in the ER-(control) spine head during the triplet stimulation, for three different choices of the pre-post spike timing difference, Δ*t*. By appropriately adjusting the plasticity thresholds *θ_D_* and *θ_P_*, our ER-spine model can show broad agreement with the experimentally reported plasticity profiles. Figure 9C shows the dependence of the Ca^2+^-CaM amplitude on the spike timing difference (Δ*t*) in the doublet (blue) and triplet (red) cases. The temporal proximity and ordering of pre-and postsynaptic inputs, together, control the NMDAR activation level, which in turn decides the maximum Ca^2+^ elevation in the spine during persistent stimulation. The corresponding synaptic weight changes (Δ*w*) induced by the STDP inputs are displayed in Figure 9D. Consistent with experimental data, the doublet protocol only induces synaptic weakening over a restricted range of Δ*t* values (Figure 9D, blue curve). On the other hand, the triplet protocol induces potentiation over an ∼35 ms window of positive Δ*t* values, flanked by two ∼40 ms windows of depression, one for Δ*t* < 0 and a second for causal pre/post pairings with longer time differences (Δ*t* ≳ +35 ms) (Figure 9D, red curve).

How does ER modulate spine Ca^2+^ signaling during STDP induction? Figures 9A-10D compare the activity-driven responses of an ER+ spine (with different IP_3_R cluster sizes) with the ER-reference spine, as a function of the spike timing difference. Release of ER store Ca^2+^ augments both the Ca^2+^ (Figure 10A) and Ca^2+^-CaM (Figure 10B) in the spine head, and this excess response in the ER+ spine relative to the ER-spine indirectly depends on the spike timing difference, which regulates the NMDAR-mediated Ca^2+^ elevation in the spine (Figures 9C and 10D). This inverse dependence of the ICCR contribution on NMDAR activation level is reflected in the NMDAR and IP_3_R Ca^2+^ current profiles, shown in Fig. 10E. Stronger NMDAR activation, particularly at small positive spike timing differences (0 < Δ*t* ≲ +40 ms), is associated with reduced ICCR, which follows from the Ca^2+^-dependent inhibition of IP_3_ receptors at sustained Ca^2+^ levels above μ0.3 μM. These trends broadly agree with the results from our previous simulations of rate-based plasticity (Fig. 6).

**Figure 9.**
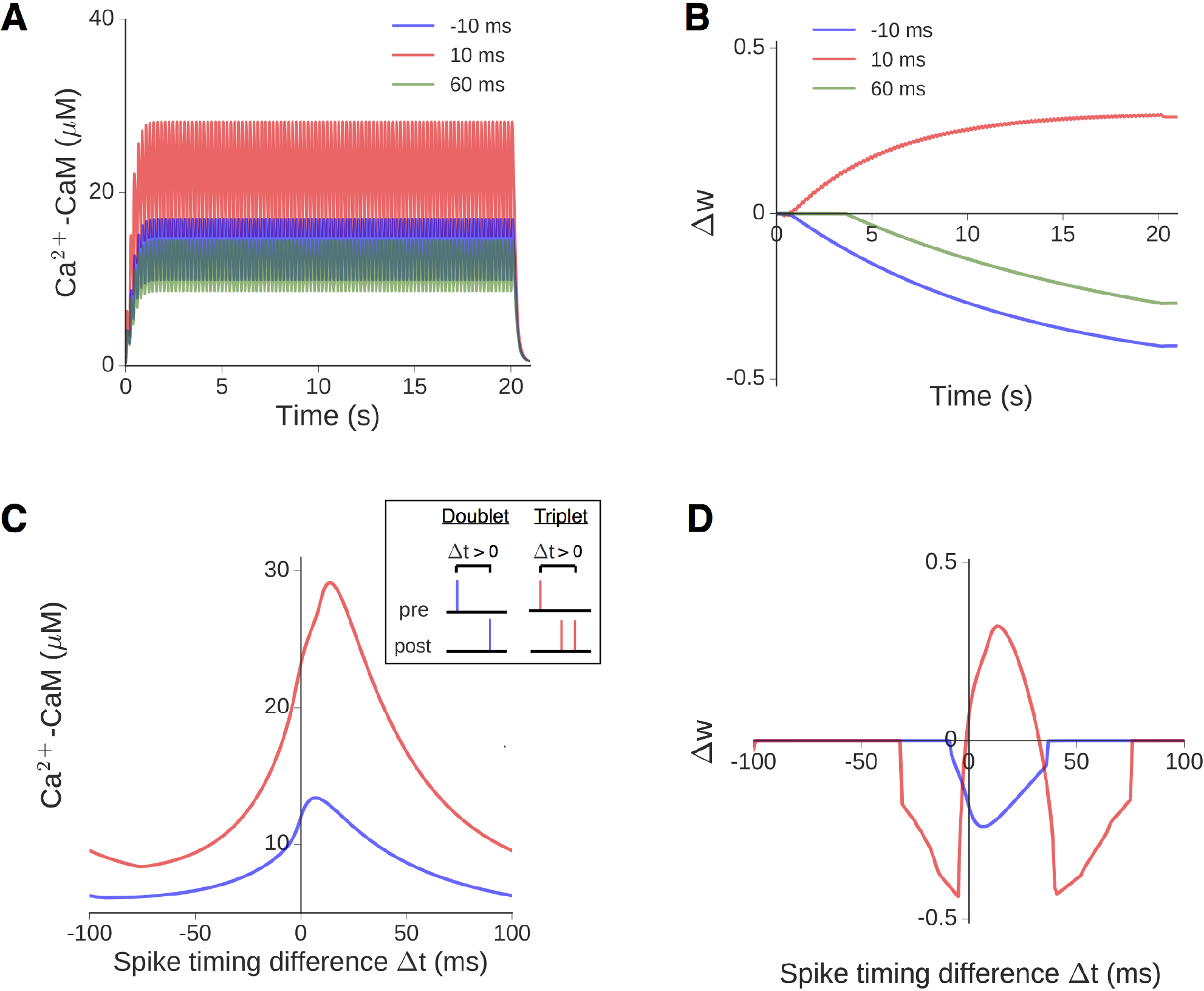
Modeling spike timing-dependent plasticity (STDP) at the ER-spine. **A,B**, Illustration of the time course of CaM activation (A) and plasticity induction (B) at the reference ER-spine in response to 100 spike triplets (1 EPSP + 2 BAP) at 5 Hz, for three different spike timing differences (Δ*t*): −10, +10 and +60 ms. Plasticity parameters have been chosen so as to yield depression at-10 and +60 ms, and potentiation at +10 ms, consistent with previous measurements^60^. **C**, Maximum CaM activation attained during prolonged paired stimulation as a function of Δ*t*, when every synaptic input is paired with either 1 (blue) or 2 (red) BAPs. The inset shows details about the STDP stimulus pattern (doublet v/s triplet pairing, and convention for positive Δ*t*). **D**, Total weight change at the end of the stimulation, plotted as a function of At for the doublet (blue) and triplet (red) spike pairings. No potentiation is induced in the former case. Parameters same as in B, in order to have overall consistency with experimental profiles.

**Figure 10.**
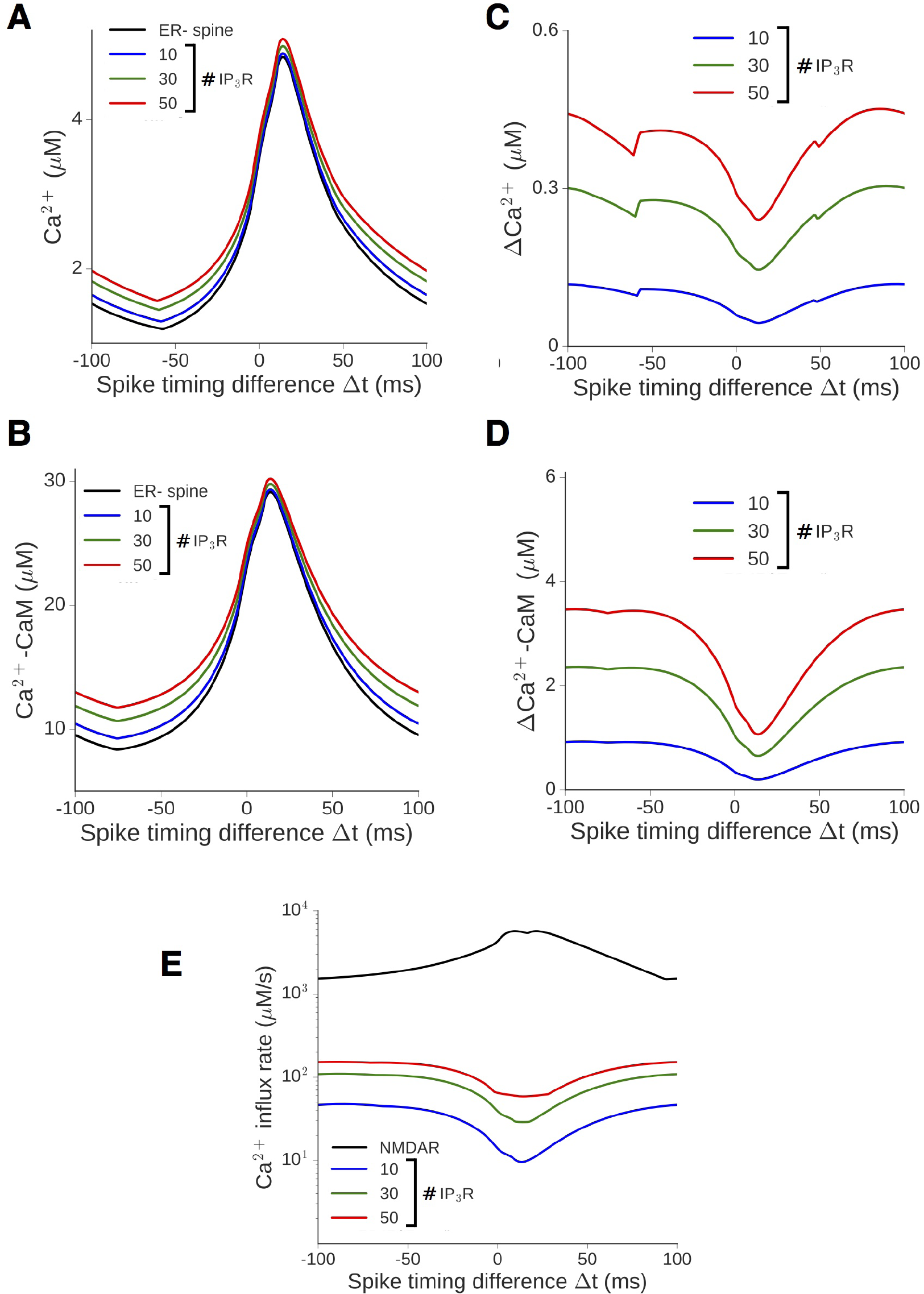
Differential Ca^2+^/CaM response in the presence of an ER depends on NMDAR activation regulated by the spike timing difference. **A,B**, The amplitude of Ca^2+^ (A) and CaM activation (B) in the ER-spine (black) versus ER+ spine for different IP_3_R numbers (colored curves), during presentation of 100 spike triplet pairings at 5 Hz over a range of Δ*t*. **C,D**, The corresponding excess Ca^2+^ (A) and CaM (B) responses in the ER+ spine relative to the ER-control. **E**, Maximum rate of Ca^2+^ influx into the ER+ spine cytosol through NMDAR channels (black) and different numbers of IP_3_R during paired stimulation over a range of At values. (All results for Δ*Ca_EPSP_* = 0.2 μM.)

Transient elevation of Ca^2+^/CaM levels drives the induction of plasticity, governed by Eq. 10. The temporal profiles of activated CaM and corresponding plasticity outcomes in the ER-/ER+ spines are illustrated for two representative spike timing differences in Figure 11. For Δ*t* = −35 ms, the amplitude of active CaM in the ER-spine lies just below the threshold for the induction of synaptic depression (Figures 9A and 11B, black curves). The additional release of ER store Ca^2+^ increases the total CaM activation above the LTD threshold, inducing strong synaptic weakening (Δ*w* < 0) in the presence of ER (Figures 9A and 11B, colored curves). When Δ*t* = +10 ms, ICCR makes a relatively modest contribution to the total Ca^2+^ response in the spine head (Fig. 11C), and this yields a small net enhancement of synaptic strengthening compared to the ER-spine (Fig. 11D). We have simulated STDP inputs to our synaptic model over the full range of allowed spike timing differences (−100 ms ≤ Δ*t* ≤ +100 ms), and the plasticity profiles obtained for the triplet and doublet protocols are shown in Figs. 12A and 12B, respectively. Ca^2+^ release from spine ER modulates the overall STDP curve for triplet inputs (Fig. 12A), and synaptic weakening is elicited over a broader range of spike timing differences compared to the ER-spine. Consistent with the reduced contribution of ICCR at small positive Δt (Fig. 10), presence of ER introduces relatively less broadening of the LTP induction window. In the case of doublet inputs (Fig. 12B), ICCR extends the window for induction of synaptic depression over a broader range of spike timing differences relative to the ER-spine, the extent of which scales with the number of IP_3_R present.

**Figure 11.**
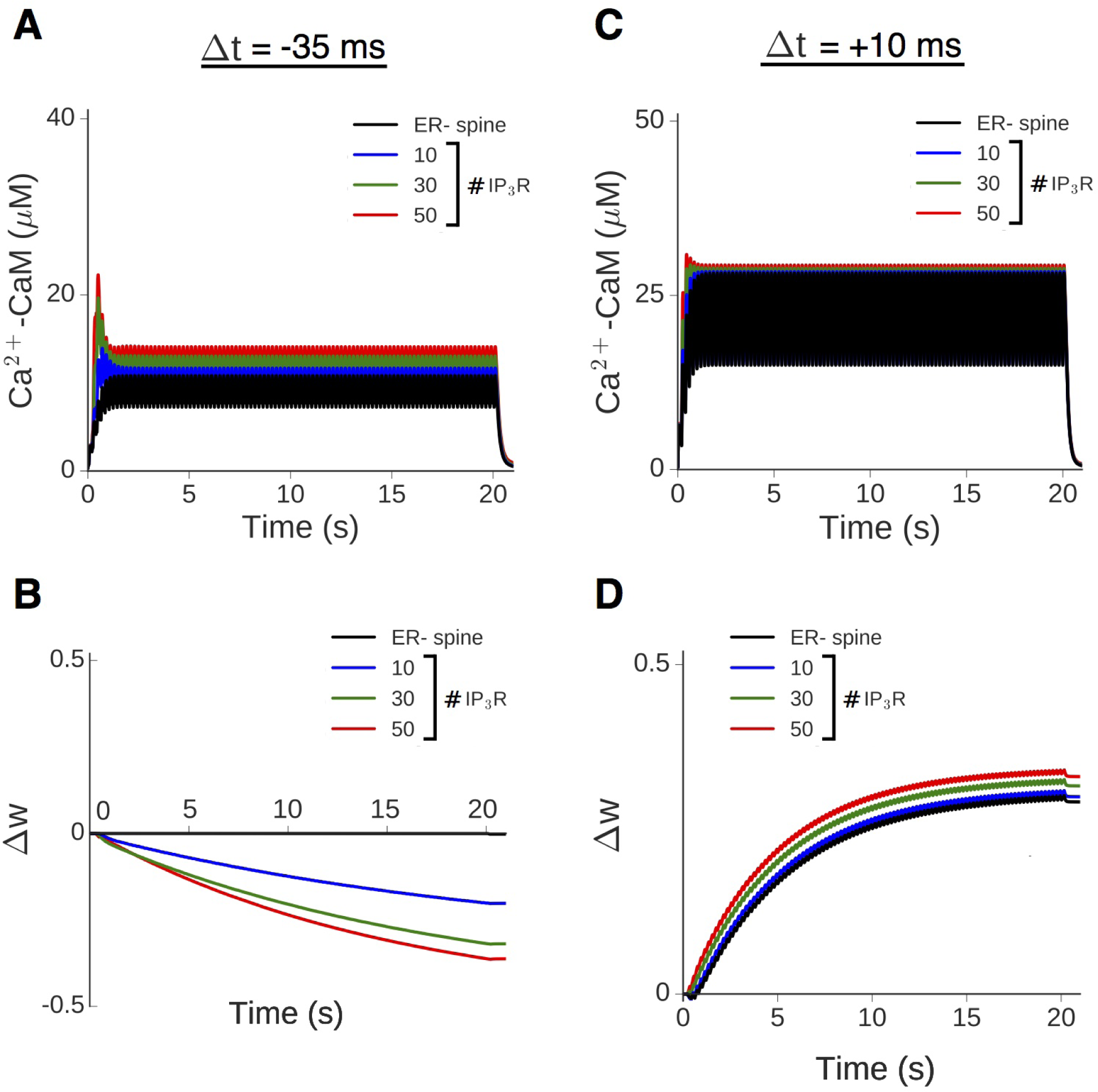
Spine ICCR modulates the induction of STDP. **A,B**, Time course of the spine CaM activation (A) and plasticity variable Δ*w* (B) in response to triplet pairing stimulation with Δ*t* = −35 ms. Results shown for the ER-control spine (black) and ER+ spine with different IP_3_R numbers (colored curves). **C,D**, Results obtained for Δ*t* = +10 ms. (Parameters here are same as in Fig. 10, with no plasticity induced in the ER-spine at Δ*t* = −35 ms.)

The differential modulation of the depression and potentiation windows suggested by the above data is quantified in terms of the relative change in widths of the LTD and LTP windows (Δ_*D*_ and Δ_*P*_) in the presence of ER. Repeated random sampling of the STDP thresholds Δ*t_D_*/Δ*t_P_* for the ER-spine (from ±10 ms windows centered on Δ*t_D_* = −35 ms and Δ*t_P_* = +35 ms) yields a distribution of possible (Δ_*D*_, Δ_*P*_) pairs. The results (aggregate of 5000 samples, 10-50 IP_3_Rs) are represented by separate heat maps for the triplet (Fig. 12C) and doublet (Fig. 12D) input patterns. The overall distribution, which, by and large, is confined to the lower triangle (Δ_*D*_ > Δ_*P*_), indicates that the contribution of ICCR can potentially extend the LTD window by several tens of ms, whereas its effect on the LTP window is comparatively less. Taken together, Figs. 12C and 12D demonstrate the relative broadening of the LTD window in the presence of ICCR which is fairly robust to variation of the thresholds and the IP_3_R cluster size. In summary, we find that mGluR-mediated Ca^2+^ release from spine ER during correlated activation of pre-and postsynaptic neurons promotes the induction of synaptic depression over a broader range of temporal activation patterns (Δ*t*), with relatively less influence on LTP induction.

**Figure 12.**
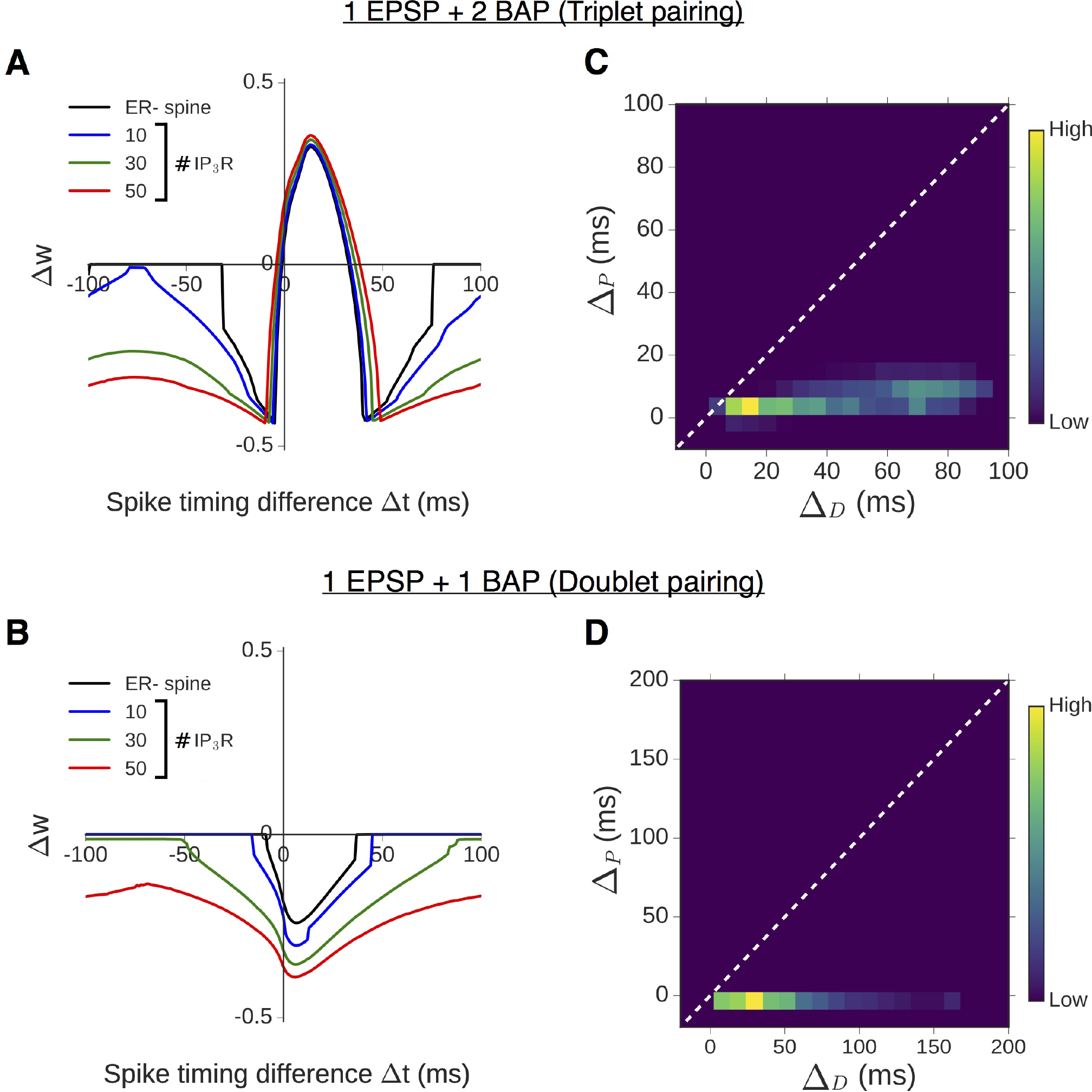
NMDAR-dependent contribution of ICCR to spine Ca^2+^ response modulates STDP profiles. **A,B**, Comparison of the plasticity profiles (total weight change Δ*w*) obtained in the absence of ER (black) and with different levels of ICCR (colored curves) in response to 100 pairings of spike triplets (A) and doublets (B). **C,D**, Quantification of relative changes in LTD and LTP window widths when ICCR is present, for the triplet (C) and doublet (D) stimulation protocols. Heat maps represent 2D distributions of (Δ_*D*_, Δ_*P*_) values obtained from random sampling (5000 times) of the STDP thresholds Δ*t_D_* and Δ*t_p_* in the ER-control spine from 20 ms windows centered on-35 and + 35 ms, respectively, and with 10-50 IP_3_R.

### Differential contribution of ICCR to LTD and LTP induction is a general consequence of IP_3_ receptor kinetics

The foregoing analysis of our detailed synaptic model highlights the potential contribution of ICCR to activity-driven Ca^2+^signaling, and its regulation by NMDAR-mediated Ca^2+^ entry into the spine. The NMDAR conductance, a key contributor to Ca^2+^, is seen to have a wide distribution in pyramidal neuron synapses^61^. To assess the robustness of our model predictions, we have systematically investigated the metaplasticity introduced by ER for a physiologically plausible range of amplitudes of synaptically evoked Ca^2+^ (Δ*Ca_EPSP_* = 0.1-1 μM) by tuning the NMDAR conductance parameter g_*N*_. g_*N*_ sets the scale for the dynamic range of NMDAR-mediated Ca^2+^ amplitudes elicited by plasticity-inducing stimuli, thereby controlling the relative contribution of ICCR to spine Ca^2+^ signaling.

Figs. 13A and 13B summarize the differential responses in the ER+ spine (for N_*R*_ = 30 IP_3_RS) relative to the ER-(control) spine during trains of synaptic (SC) input at different rates. The heat map in Fig. 13A represents the excess Ca^2+^-CaM amplitude as a function of the input frequency (horizontal axis) and g_N_ (vertical axis). ER contributes robustly to spine Ca^2+^ elevation at low input frequencies (*f* ≲ 5 Hz). The contribution of ICCR steadily declines at higher input rates due to Ca^2+^-dependent inhibition of IP_3_ receptors, and this frequency-dependent suppression of the difference between the ER-and ER+ spines is more pronounced at higher Δ*Ca_EPSP_*. The corresponding results for the differential induction of plasticity in the ER+ spine are displayed as a heat map in Fig. 13B. For each g_*N*_, the plasticity thresholds and *θ_D_* and *θ_P_* have been adjusted to have *f_D_* = 1 Hz and *f_P_* = 15 Hz in the ER-spine, and the excess plasticity (ΔΔ*w*) in the presence of ER has been estimated for every input frequency. Fig. 13B indicates that ICCR robustly augments NMDAR-mediated Ca^2+^ responses to facilitate the induction of synaptic depression at low input frequencies (*f* ≲ 1 Hz). Due to the inhibition of ICCR with increasing NMDAR activation (Fig. 13A), there is little difference between the plasticity curves for the ER-and ER+ spines at higher input frequencies. Except in a limited range of small g_*N*_ values (Δ*Ca_EPSP_* ≲ 0.15 μM), ICCR is strongly suppressed at stimulation rates above ~10 Hz, and the LTP threshold (*f_P_*) in the ER+ spine remains nearly unchanged relative to the ER-spine.

**Figure 13.**
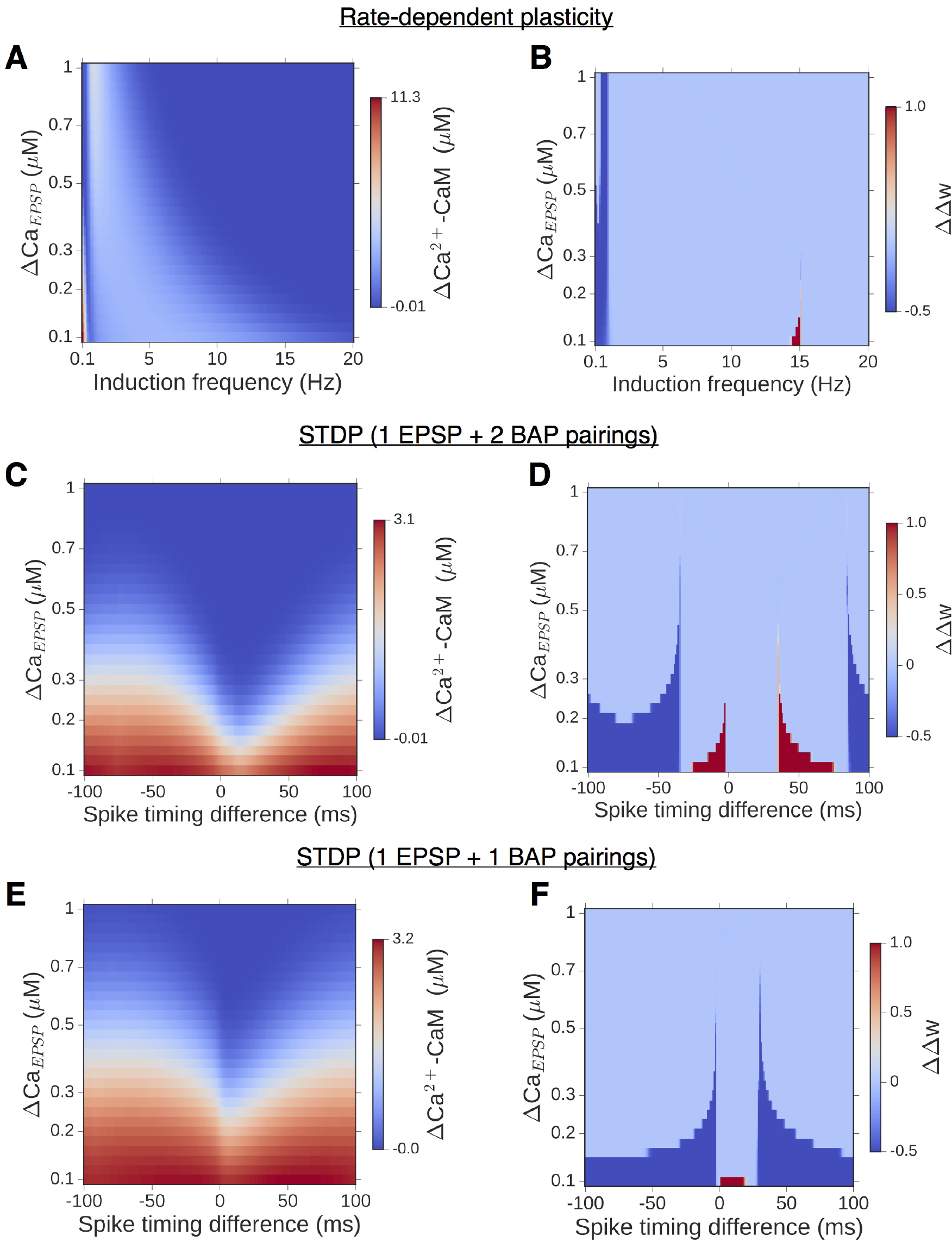
Differential enhancement of LTD and LTP by spine ER is a general consequence of ICCR kinetics. **A,B**, Dependence of the differential CaM activation (A) and differential synaptic weight change (B) at the end of stimulation on NMDAR conductance (Δ*Ca_EPSP_*) and the synaptic stimulation rate. **C-F**, Parameter sensitivity in the case of STDP inputs for the triplet (C,D) and doublet (E,F) spike pairing stimuli. (All results for comparison between ER-control spine and an equivalent ER+ spine with 30 IP_3_R; plasticity thresholds appropriately adjusted to give (in the ER-spine) *f_D_/f_P_* = 1 Hz/15 Hz for rate-dependent plasticity, and, separately, STDP for-35 ms < Δ*t* < 70 ms.)

We have similarly characterized the differential responses and plasticity outcomes in the ER+ spine during STDP input patterns (Figs. 13C-13F). For each g_*N*_, the plasticity thresholds *θ_D_* and *θ_P_* in our model were adjusted so as to yield (for spike triplets) an LTD window for −35 ms < Δ*t* < 0 ms and a potentiation window for 0 ms < Δ*t* <35 ms in the ER-spine. Fig. 13C summarizes the dependence of the excess Ca^2+^-CaM response in the ER+ spine (with N_*R*_ = 30 IP_3_Rs) on g_*N*_ and the spike timing difference (Δ*t*) in the triplet case, showing a general reduction in the ICCR-mediated enhancement of spine Ca^2+^ signals with increasing Δ*Ca_EPSP_*. The corresponding differences in the plasticity outcome (ΔΔ*w*, Fig. 13D) show significant broadening of the window of LTD induction over a fairly wide range of g_*N*_ values. The ER-associated broadening of the LTP window is restricted to a comparatively narrow range of g_*N*_ values and spike timing differences. The differential CaM response and plasticity outcome in the doublet case (Figs. 13E and 13F, respectively) similarly highlight a general broadening of the window of spike timing differences eliciting synaptic depression, the extent of which scales inversely with Δ*Ca_EPSP_*.

As the above results have been obtained for a specific choice of the plasticity thresholds and IP_3_R cluster size, we repeated these comparisons with variable number of IP_3_ receptors (N_*R*_ = 10-50) and plasticity thresholds, for both frequency-dependent plasticity and STDP (Methods). For each set of parameter values, the overall modulation of the NMDAR-only plasticity curve by ICCR was quantified in terms of the shifts in the LTD and LTP thresholds for rate-dependent plasticity, and changes in the LTD/LTP window widths in the case of STDP. The summary statistics (Fig. S2) provide a sense of the variability in model outcomes, and taken together, suggest graded modulation of NMDAR-based bidirectional plasticity by ICCR, with selective enhancement of LTD induction. In sum, our model simulations over realistic parameter ranges provide support for an ER-associated form of synaptic metaplasticity on the level of individual spines, the nature of which is regulated by mGluR-IP_3_ signaling in concert with the temporal profile of NMDAR activation.

## Discussion

Dendritic spines are specialized structures that facilitate spatially restricted biochemical signaling and enable input-specific, Hebbian-type synaptic changes mediated by NMDA receptor^62, 63^. In the present study, we have systematically investigated, with mathematical modeling, how the local reorganization of ER may modulate Ca^2+^-driven plasticity on the scale of individual excitatory CA1 synapses. Results from our model simulations of rate-based plasticity and spike timing-dependent plasticity, taken together, suggest that the presence of ER selectively enhances the propensity for LTD induction with a relatively diminished effect on LTP induction. Targeting of ER to larger spine heads can thus mediate a form of metaplasticity at stronger synapses that specifically relies on ER Ca^2+^ handling and metabotropic glutamatergic signaling. The graded contribution of the IP_3_-sensitive ER store to spine Ca^2+^ elevation as a function of NMDAR activation that we have characterized here offers a synapse-specific mechanism to differentially modulate the windows for depression/depotentiation and potentiation, yielding a net enhancement of activity-driven synaptic depression (Fig. 14). In light of the observed association of ER with more potent synapses^30^ and the dynamic acquisition of ER accompanying spine enlargemen^36, 38^, we propose a regulatory role for ER as a “braking” mechanism, that may temper the propensity for further strengthening at the potentiated synapses. Our findings thus support a novel interpretation for spine ER in adjusting the plasticity profile of a synapse on an as-needed basis, which may contribute to keeping saturation at bay, and maintaining the distribution of synaptic strengths in a useful dynamic range supporting memory storage and optimal neural network functio^2, 6, 14^.

**Figure 14.**
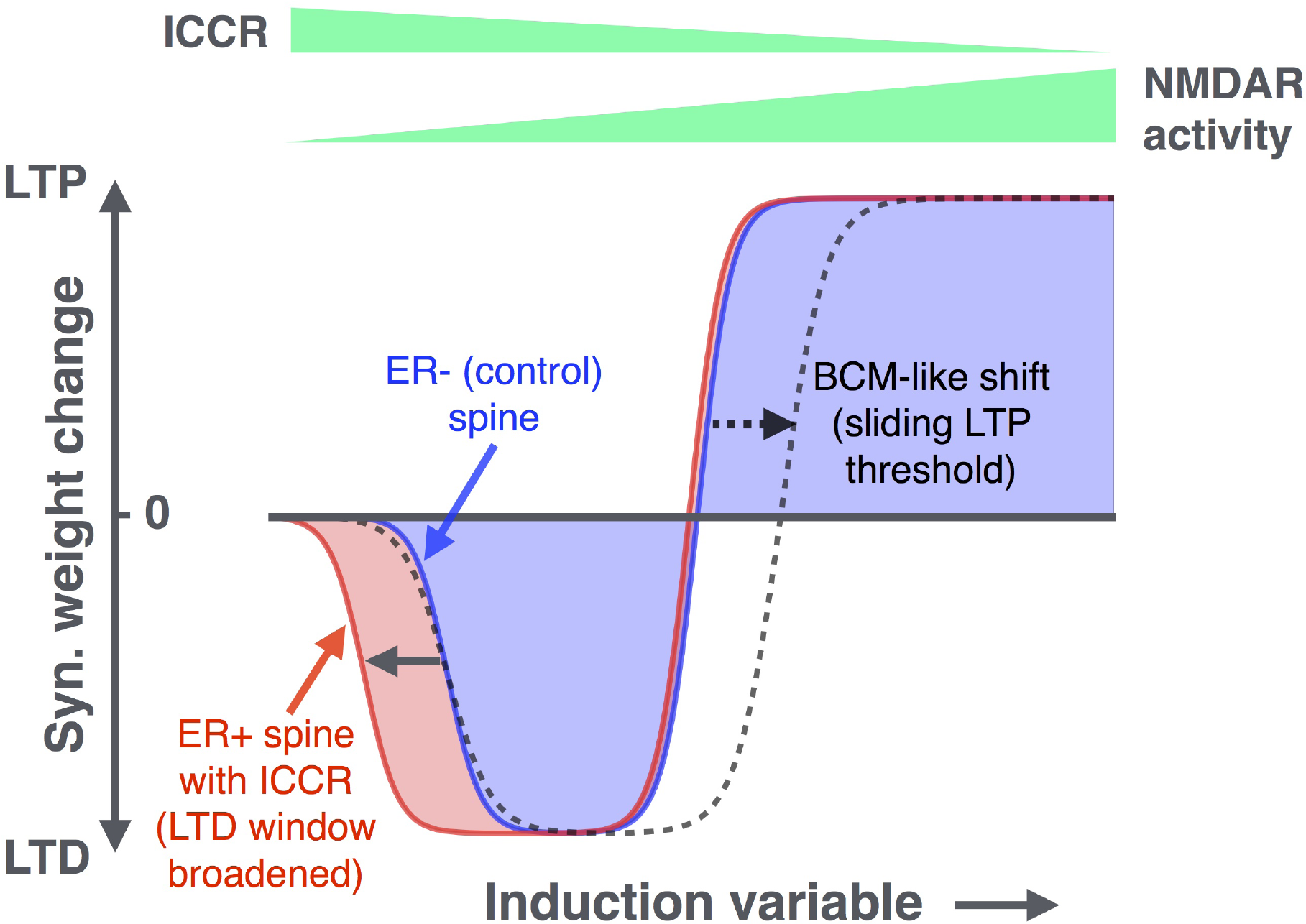
ER Ca^2+^ store introduces a novel form of synaptic metaplasticity at individual dendritic spines. Results from our analysis of frequency-and spike timing-dependent plasticity are summarized by a comparison of the plasticity profiles for the reference ER-spine (blue) and a spine with ER (red). The induction variable (which can stand for the synaptic input frequency *f*, or spike timing difference Δ*t*) controls the activation of both NMDAR and ICCR, these dependencies being represented by tapering bars at the top. Also shown for comparison is the modified plasticity curve arising from a BCM-like sliding LTP threshold (dashed curve).

Blocking synaptic strengthening in the hippocampal formation induces amnesia^64^ and suggests that synapse-specific hippocampal LTP is essential for initiation of experience-dependent learning. This locus is shifted subsequently and cortical neural circuits take over memory consolidation and retrieval after initial memory formation is complete^64, 65.^ Spines having undergone experience-dependent modifications can quickly saturate (runaway effect of LTP) and no longer participate in ongoing learning. Adaptability of the plasticity profile may promote reuse of strong synapses, enabling saturated synapses to be brought back into the ‘game’. The form of metaplasticity elucidated here is specific to individual CA1 spines, and is likely triggered by the rapid local remodeling of ER that may be a consequence of strong prior synaptic activation^36^. Our work is of particular relevance to an understanding of the mechanisms in place to overcome possible storage limitations in the CA1 fields, given that ER is most likely to be found associated with stronger synapses at the larger CA1 spines. The selective enhancement of synaptic weakening in the presence of ER (Fig. 14) has a direct implication for long-term stability of the stronger synapses: under “natural” conditions^66^, background synaptic activation may promote a slow resetting of potentiated synapses at the ER-bearing spines; viewed another way, association with ER may be seen to result in an effective “slowing down” of cumulative synaptic strengthening over time. We note that these effects are distinct from a BCM-like shift of the LTP threshold^7^, though at higher SERCA activity levels, ER may introduce such a shift in addition to promoting LTD induction (SI and Fig. S6).

Computational models of Ca^2+^-based plasticity normally attempt to link NMDAR-mediated Ca^2+^ entry to downstream signaling events at the postsynaptic locus, and these have provided a meaningful account of the general phenomenolgy of long-term plasticity in the hippocampus and neocorte^49-53, 67^. From a neurobiological point of view, though, it is important to go beyond generic descriptions and consider whether local heterogeneities in microphysiology on the synapse level could introduce functional differences between individual synaptic connections in a population. The role of spine ER in introducing synapse-specific functional differences in Ca^2+^ signaling may not be clearly discernible from macroscopic measurements of synaptic plasticity properties, or by lumping together and averaging over experimental data from many synaptic contacts only a few of which may be associated with ER-bearing spines. This situation in hippocampal pyramidal neurons may be contrasted with the case of spines associated with cerebellar parallel fiber to Purkinje cell synapses, which are more homogeneous in terms of the presence of ER^68^ and where localized Ca^2+^ release from stores is known to be central to the induction of long-lasting plasticit^69-71^.

This last aspect, in particular, underlines the utility of a physiogically realistic *in silico* model which enables systematic characterization of the role of ER in Ca^2+^ signaling at individual spines. The framework presented here is one of the first to weave a quantitative description of mGluR-and IP_3_-regulated signaling into a detailed model of spine Ca^2+^ dynamics driven by NMDAR activation. Delineation of the role of ER Ca^2+^ handling in a spine requires considering mGluR-IP_3_ signaling in the backdrop of synaptic NMDAR activation, as NMDAR-gated Ca^2+^ entry can regulate several aspects of this signaling, including PLCβ activity, IP_3_ turnover, and IP_3_R gating, with dynamic Ca^2+^ feedback from ICCR further adding to the overall system complexity (Fig. 1C). We have calibrated the parameters in our model of ICCR to be consistent with salient features of experimentally reported Ca^2+^ responses in ER-containing CA1 spines to unitary synaptic events^32^, particularly the lag (few hundred ms) in Ca^2+^ release from ER following release of glutamate into the synaptic cleft. Our analysis builds on these observations as we examine the contribution of ER to the spine Ca^2+^ response, and its subsequent shaping of plasticity, in the context of neural activity patterns that mimic the experimental induction of early LTP/LTD.

The response of an ER+ spine in our model to low-frequency glutamate pulses, in particular, highlights the interaction of multiple timescales associated with IP_3_ degradation, recovery of IP_3_R from Ca^2+^-dependent inactivation, and the input frequency in shaping the profile of ICCR. Our model yields a non-monotonic rate dependence of the ER Ca^2+^ contribution at low input frequencies (*f* ≲ 5 Hz) (Fig. 4). As noted before, this arises from a balance between the contrasting effects of the input rate on (1) recovery of IP_3_R from inactivation between inputs (which is lesser, the more closely spaced the inputs are), and (2) the availability of IP_3_, which increases with input rate and promotes the opening of IP_3_R channels (Fig. S3A). Mirroring the profile of ER Ca^2+^ contribution, our model also predicts a non-monotonic rate dependence for the delay in ICCR following glutamate release (Fig. S3B), which initially increases before decreasing with the input frequency as ICCR gradually synchronizes with the NMDAR-mediated Ca^2+^ transient. The foregoing properties of ICCR dynamics inferred from our numerical simulations are potentially informative readouts of the model, which could help to validate the overall kinetics and constrain key model parameters by comparison with future experimental measurements.

The contribution of ICCR to spine Ca^2+^ signaling can be broadly understood in terms of NMDAR-mediated spine Ca^2+^regulation and the kinetics of IP_3_R with its characteristic bell-shaped dependence on the cytosolic Ca^2+^ level. With repeated low-frequency stimulation, there is no sustained build-up of Ca^2+^ evoked by the sequence of glutamate pulses. Every synaptic input triggers transient mGluR activation, evoking a short burst of Ca^2+^ release from ER which is initiated by fast activation (m_1_ and m_2_) of IP_3_R flux, followed by the slower Ca^2+^-dependent inactivation (*h*) of the IP_3_R which brings the Ca^2+^ level down. This allows the IP_3_R to recover from inactivation between successive inputs when Ca^2+^ has decayed back to near-resting levels, and explains the robust augmentation of the NMDAR-mediated Ca^2+^ signal by ICCR at low stimulation frequencies (Fig. 4; also Figs. S4A-S4D). At higher input frequencies, NMDAR-gated Ca^2+^ transients triggered by successive inputs add up, and there is sustained elevation of Ca^2+^ in the spine (see, e.g., Figs. S4E and S4F), the scale of which is set by the input frequency *f*. As Ca^2+^ is continually maintained at a high level, the Ca^2+^-dependent inactivation variable h stays small, and keeps the IP_3_R persistently inhibited as long as the stimulation is present, despite the concurrent activation of mGluR and availability of IP_3_. This underlies the progressive, frequency-dependent suppression of Ca^2+^ flux through IP_3_ receptors at higher input frequencies (Fig. 6), and implies a steadily diminishing role for ICCR as LTD switches to LTP according to the Ca^2+^-based plasticity model governing bidirectional synaptic strength changes (Fig. 8).

An analogous situation arises during trains of pre-and postsynaptic spikes at a constant rate (in the theta band), mimicking the experimental induction of spike timing-dependent plasticity. The activation of NMDA receptors in this case is controlled by the relative timing of pre-and postsynaptic firing (Δt) on the millisecond scale. The amplitude of the sustained Ca^2+^ elevation driven by NMDA receptor activity is thus a function of the spike timing difference, and it directly controls the level of inhibition (*h*) of the IP_3_R (Eq. 7). This accounts for the spike timing dependence of the relative contribution of ICCR to the total spine Ca^2+^ signal in Fig. 10, and the overall enhancement of synaptic depression at the ER+ spine (Fig. 12).

Regulation of synapse strength is acknowledged to be a complex process, likely involving the coordinated action of several mechanisms that may act over a wide range of time and spatial scales to modulate the rate and direction of ongoing activity-dependent plasticity. Previously proposed mechanisms include intrinsic plasticity of membrane excitabilit^13, 72^, compensatory scaling of spine volume to balance synaptic potentiatio^73, 74^, alterations in the number and/or subunit composition of postsynaptic NMDA^12, 75^, global synaptic scaling via glial signaling^76^, etc. The present study linking ER to local modulation of Ca^2+^-based plasticity adds to the repertoire of dynamically regulated biophysical mechanisms that may be in place to control plasticity and function at excitatory hippocampal synapse^5, 6^.

Although several lines of evidence link group I mGluR activation and ER Ca^2+^ stores to long-term depression at hippocampal synapses, the involvement of ER, or more generally of mGluR signaling, in synaptic potentiation is less clear. Results from several pharmacological and knockout studies collectively do not implicate an essential role for group I mGluR in LTP or AMPAR-mediated synaptic transmission in the CA1 region^77^. Mutant mice lacking a function G-protein associated with group I mGluR were found to be deficient in hippocampal LTD induced by low-frequency stimulation, but exhibited intact LTP in response to tetanic inputs^43^. Some studies on mGluR5 knockout mice reported reduced LTP in CA1 neuron^78, 79^, but this was shown to involve selective reduction of NMDAR function on both the induction and expression levels, with no effect on the AMPAR component of LTP. Further, there is no direct evidence that the above interaction relies on Ca^2+^ release from stores, or that it is synapse-specific and restricted to spines containing ER. As noted previously, LTP studies usually examine plasticity on the coarse-grained level and not at individual synapses; thus, they are of limited utility in addressing the differences in local Ca^2+^ signaling that might arise between ER-and ER+ spines during LTP induction. Moreover, the concurrent stimulation of multiple synaptic contacts (as in experimental induction of plasticity) may evoke Ca^2+^ release from ER in the dendritic body as well^80^; this could activate signaling pathways distinct from those associated with early LTP in spines, and trigger plasticity at a large number of synapses (spread out over a wider dendritic region), independent of their association with ER-bearing spines. The present study is restricted to quantifying the contribution of ER (IP_3_-sensitive stores) to spatially confined Ca^2+^ dynamics in spines, and it is in this specific context that our analysis predicts decreasing ER contribution with stronger synaptic (NMDAR) activation due to the Ca^2+^-dependent suppression of IP_3_R activity. We note that an early experimental study of STDP at CA3-CA1 synapses^58^ makes a similar suggestion about the suppression of IP_3_R activity during LTP (but not LTD) induction, as a possible explanation for the absence of associated heterosynaptic plasticity mediated by regenerative Ca^2+^ release (Ca^2+^ waves) from IP_3_-sensitive stores.

To conclude, we have presented a detailed modeling study, leveraging previous experimental observations, to characterize the local modulation of Ca^2+^ signals and plasticity at hipocampal dendritic spines by ER. Our analysis lends support to the view that targeting of ER to individual spines could regulate their potential for subsequent plasticit^32, 36, 38^, providing a local mechanism to alter the rules by which activity patterns are transformed into synaptic efficacy changes at individual synapses. The suggestion of a synapse-specific modulatory role of ER made here adds a new dimension to earlier work on metaplasticity mediated by metabotropic receptor^14, 81-83^. Incidentally, a role for store Ca^2+^ release was also recently implicated in synaptic homeostasis mediated by spontaneous miniature EPSPs^84^. The involvement of ER (as a Ca^2+^ store) in biochemical signaling likely extends to other aspects of dendrite function, such as in mediating some forms of heterosynaptic plasticity^58^, long range (synapse-to-nucleus) signaling involved in activity-dependent transcriptional regulation^85^, spatiotemporal integration of synaptic inputs^86^, and may also be relevant for mechanistic understanding of cognitive disorders such as Alzheimer’s and Fragile X syndrome which have recently been linked to aberrant mGluR signaling and Ca^2+^ store dysregulation at hippocampal synapse^87-90^. The approach presented here provides a framework for future modeling studies aimed at investigating these varied questions. On a more general note, the potential contribution of ER to microscale signaling elucidated here highlights the need to move beyond “average” accounts of synaptic signaling, and consider the implications of local physiological differences between synapses for their plasticity and function.

## Methods

We implement a deterministic, single compartment model of a spine head on a hippocampal CA1 apical dendrite, and characterize the calcium (Ca^2+^) elevation and early long-term plasticity driven by synaptic activation in the presence of an ER/spine apparatus (Fig. 1). The spine head is modeled as a sphere with fixed volume V_*spine*_ = 0.06 μm^3^, which approximates an average ER-bearing spine head (ER+ spine) found experimentally^32^. While ER+ spines are on average larger than spines lacking an ER, our canonical spine head lies well within the reported dynamic range of spine sizes/synaptic strength^32, 91^, and thus may be considered potentially capable of undergoing plasticity in both directions (strengthening as well as weakening of synaptic efficacy). ER extending into spines is contiguous with the dendritic ER^39^, and is typically found to occupy only a small fraction of the spine volume (≲ 5%)^29^; we assume an ER-to-spine head volume ratio of 0.1. The spine head is assumed to be electrically coupled to its parent dendritic shaft by a thin “neck”, modeled as a passive electrical resistance of *R_C_* = 100 MΩ^92^. Diffusive coupling of Ca^2+^ between the spine and dendrite is ignored, in accordance with Ca^2+^ measurements at mushroom spine^91, 93^.

The various biophysical components comprising our model that collectively regulate the electrical and Ca^2+^ dynamics in the spine are described below. All model parameters and molecular concentrations used in our simulations are listed in Tables S1-S4.

### Membrane voltage dynamics at the spine

The voltage at the postsynaptic membrane (u) is described by the following Hodgkin-Huxley (HH)-type ordinary differential equation:

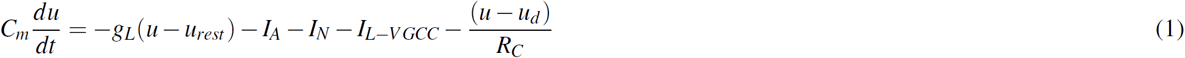

which includes contributions from a passive leak current (g_*L*_), voltage dependent AMPA receptor (AMPAR)/NMDA receptor (NMDAR)-gated currents (I_*A*_/I_*N*_) in the PSD, a high voltage-activated L-type Ca^2+^ current (I_*L-VGCC*_), and passive electrical coupling to the dendritic shaft (R_*C*_) with *u_d_* denoting the voltage of the dendritic compartment (Fig. 1B). We assume standard membrane parameters (capacitance C_*m*_ = 1 μF/cm^2^ and uniform leak conductance density *g_L_* = 0.0002 S/cm^2^), and the resting membrane potential in both the spine and parent dendrite is set to *u_rest_* =-70 mV^50^.

AMPAR and NMDAR-gated currents are assumed to be transiently activated every time a synaptic input arrives, and both are modeled with linear I-V relations (reversal potentials E_*A*_, E_*N*_ = 0 mV)^67^. The AMPAR conductance is modeled as the difference between two exponentials with a rise time constant of 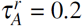 and decay time constant 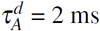^50^:

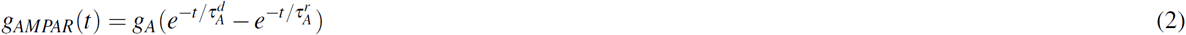

for a glutamate pulse arriving at *t* = 0. The conductance parameter g_*A*_ is fixed at 0.5 nS across all our simulations^67^. The glutamate dependence of the total NMDAR current is similarly modeled as a difference between exponentials with longer response times 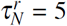 and 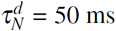)^13^; the NMDAR conductance has an additional (multiplicative) dependence on the membrane potential, describing the Mg+ block, which is modeled as a sigmoid function of the form *B*(*u*) = 1/(1 + 0.28 exp(−0.062*u*))^50^. The NMDAR conductance parameter g_*N*_ is a variable in our analysis, and adjusted to obtain different levels of spine Ca^2+^ elevation evoked by unitary synaptic input (see below). Trains of synaptic stimulation have been modeled as the sum of the above conductance waveforms^67^.

The L-type voltage-gated Ca^2+^ current is regulated by HH-type activation and inactivation gating variables, *m_u_*(*t*) and *h_u_*(*t*), respectively, whose time dependence is governed by the following equations^94^:

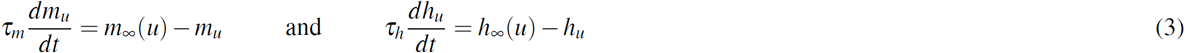

where 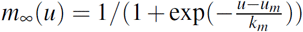 and 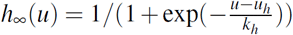, with *u_m,h_, k_m,h_* and *τ_m,h_* listed in Table S1. The total (i.e. Ca^2+^) current through the L-VGCC is described by a modified Goldman-Hodgkin-Katz (GHK) relation^95^ in order to correctly account for the large Ca^2+^ concentration gradient between the cytosol ([Ca^2+^]_*rest*_ = 50 nM) and the exterior of the cell ([Ca^2+^]_*ext*_ = 2 mM) under basal conditions. It is given by

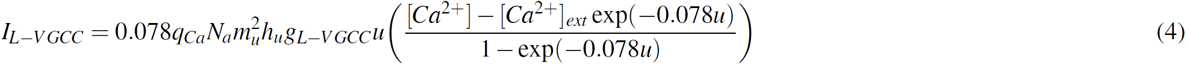

where *q_Ca_* is the electric charge per Ca^2+^ ion, N_*a*_ is the Avogadro number, and the conductance parameter g_*L-VGCC*_ is set according to the value of g_*N*_ such that the peak L-VGCC Ca^2+^ influx rate during a backpropagating action potential (BAP) is comparable to that mediated by NMDAR in response to a glutamate pulse (i.e., during an evoked postsynaptic potential (EPSP)), consistent with single spine measurements^91^.

### Calcium regulation

We assume a basal steady-state Ca^2+^ level of 50 nM in the spine head^67^. The time course of the averaged, free (i.e. unbound) Ca^2+^ concentration during synaptic activity is described by the following equation:

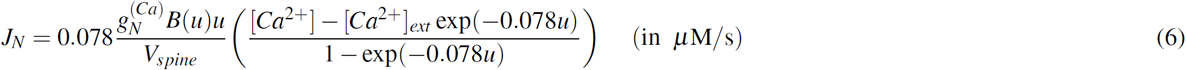

The terms J_*ER*_ represents the overall contribution of spine ER to Ca^2+^ activity (described in detail in the next two subsections), and is set to zero in the ER-spine head, which in all other respects is identical to the ER+ spine. The Ca^2+^ influx through the NMDAR channel cluster, J_*N*_, is also described by a GHK-type current term, and as in previous studies, we have assumed that the Ca^2+^ current constitutes 10% of the total NMDAR curren^50, 95^:

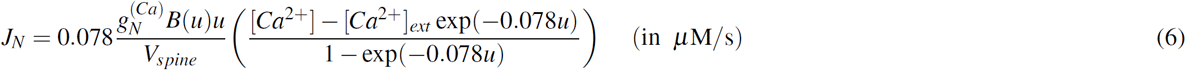

where 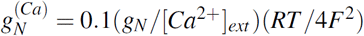, R denoting the universal gas constant (8.314 J mol^-1^ K^-1^), F = 96485 C mol^-1^being the Faraday constant, and the temperature T set to 30 °C.

The term *J_B_* encapsulates the net effect of cytosolic buffers (Table S2). Calbindin D28-k (CB) is a prominent fast-binding mobile buffer in hippocampal neurons^96^. The interaction of Ca^2+^ with CB is described by a detailed nine-state kinetic scheme^93^ (Fig. S5A). Each CB molecule carries two high-affinity and two medium-affinity binding sites for Ca^2+^. The total concentration of CB is taken to be 45 *μ*M, as measured experimentally in CA1 pyramidal neurons^96^. Besides CB, we include an endogenous “immobile” Ca^2+^-binding protein (CBP) with total concentration of 80 *μ*M, as inferred previously from a spatial reaction-diffusion model of spine Ca^2+^ transients fit to single-spine experimental data^93^. Its interaction with Ca^2+^ is described by a first-order reversible reaction (Fig. S5B). In addition, we include a slow buffer with a total concentration of 40 *μ*M, the kinetic parameters for which are set ten times slower than those for the immobile buffer^97^. We also include calmodulin (CaM) in our simulations. CaM is known to be present at a high concentration in CA1 dendritic spines^98^. Due to the slower kinetics of its interaction with Ca^2+^ compared to CB, it has little impact during short Ca^2+^ transients (single synaptic events), but is expected to make a significant contribution to regulating free Ca^2+^ levels on longer timescales during persistent stimulation. We have set the total CaM concentration to be 50 */J,*M, which represents an average of several estimates found in the literatur^67, 99, 102^. We adopted a kinetic scheme used previously^103^ to describe the reversible binding of CaM to Ca^2+^. Each CaM molecule comprises a high-affinity C-lobe and a low-affinity N-lobe, each of which can cooperatively bind up to two Ca^2+^ ions (Fig. S5C).

In addition to the Ca^2+^ currents and buffers, the Ca^2+^ level in the cytosol is also regulated by efflux mechanisms operating at the plasma membrane (denoted by the J_out_ term in Eq. 5), which set the time scale for the decay of Ca^2+^ transients in the spine head. We model efflux via plasma membrane Ca^2+^-ATPase pumps (PMCA) and sodium-calcium exchangers (NCX), both of which are approximated by Michaelis-Menten-like enzyme kinetics (Fig. S5D). Surface densities and kinetic parameters for PMCA and NCX are adopted from a previous modeling study of CA1 spines^93^. In order to balance the extrusion of Ca^2+^ and ensure that Ca^2+^ in the cytosol can be maintained at a basal level of 50 nM in the absence of stimulation, the transporter equations also include leak terms (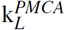 and 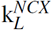), which are set so as to exactly balance the efflux rate per pump molecule when the intracellular Ca^2+^ is held at 50 nM (Table S2).

### Model for IP_3_-and Ca^2+^-induced calcium release (ICCR) from spine ER store

The intracellular Ca^2+^ store associated with spine ER/spine apparatus is a potential additional source of Ca^2+^ elevation in the spine head during synaptic activation. We model Ca^2+^ release from the ER pool evoked by the activation of G protein-coupled group 1 metabotropic glutamate receptors (mGluR) (assumed to be present on the postsynaptic membrane), subsequent production of the second messenger IP_3_, and the downstream activation of IP_3_ receptors on the ER membrane (Figs. S5E-S5G). We have adapted a previously published kinetic description of mGluR activation leading to PLCβ-mediated hydrolysis of PIP2 at cerebellar PF-PC synapses^54^, which itself borrows heavily from earlier models of metabotropic signaling proposed for generic mammalian neuron^104, 105^. Glutamate uncaging at individual CA3-CA1 synapses^32^ indicates a delay of a few hundred ms in Ca^2+^ release from the spine ER following the presentation of glutamate. A simpler (Hill function-based) model for mGluR activation and IP_3_ turnover is unable to reproduce this experimental delay, therefore, we used the detailed kinetic description, which allows for more flexibility in tuning the time course of IP_3_ production.

For the purpose of modeling mGluR-Gq activation, we simulated each glutamate pulse (synaptic input) as an alpha function with a time constant of τ_giu_ = 1 ms (glutamate is rapidly cleared from the synaptic cleft by the high density of glutamate transporters present on the surrounding astrocytic membrane^93^), and a peak concentration of 300 *μ*M at the perisynaptically located mGluR^54^ (scaled down from ~1 mM at the PSD). We asked which of the parameters in the detailed mGluR-IP_3_ model most affected the latency in IP_3_ production. The IP_3_ time course was found to be robust to variation in all except a combination of 3 parameters (a_1*b*_, a_2*b*_ and b_11_), which were scaled down appropriately (a_1*b*_ and a_2*b*_ by a factor of 50 and bn by a factor of 4) to introduce a few hundred ms delay in the peak of the IP_3_ response following the initial input. All other parameter values governing Gq-mediated PLC activation and IP_3_ turnover were taken to be the same as in the original study^54^ (Table S3).

The gating of IP_3_ receptors is modeled with a reduced kinetic scheme adopted from a previous computational study of ER Ca^2+^ dynamics in neuroblastoma cells^106^. IP_3_R activation is assumed to depend on the instantaneous levels of IP_3_ and Ca^2+^, modeled by the activation variables *m*_1_ = [IP_3_]/(d_1_ + [IP_3_]) and *m*_2_ = [Ca^2+^]/(d_5_ + [Ca^2+^]), with half-activation constants d_1_ = 0.8 μM (lower affinity compared to *in vitro* measurements^107^) and d_5_ = 0.3 μM. IP_3_R are inactivated at higher calcium levels, which is accounted for by the slower kinetics of a third gating variable, *h*, given by the following equation:

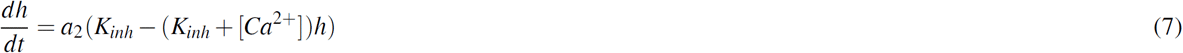

with the dissociation constant for inhibition, K_*inh*_, set equal to 0.2 μM. The Ca^2+^ influx through the IP_3_R-gated channels is given by

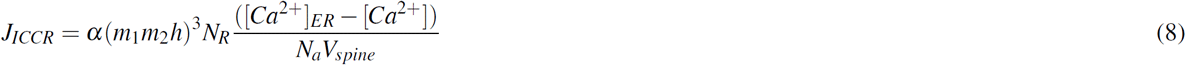

where N_*R*_ is the cluster size of the IP_3_R-gated channels present in the spine head (estimated to be in the range of a few ten^54, 108, 109)^, and *α* sets the magnitude of the Ca^2+^ current through an open channel for unit concentration difference across the ER membrane. Single channel measurements suggest a type 3 IP_3_R open channel current of μ0.15 pA at a Ca^2+^ concentration difference of 0.5 mM^110^. The Ca^2+^ concentration in the ER lumen is assumed to remain constant in our simulations, and we use a value of [Ca^2+^]_*ER*_ = 250 μM throughout, which represents an average of several estimates in the literatur^28, 111, 112^.

The ER can also contribute to cytosolic Ca^2+^ efflux through the sarco-endoplasmic reticulum ATPase (SERCA) transporters present on the ER surface, which maintain the steep concentration gradient (~10^3^-10^4^) across the ER membrane. The SERCA activity is assumed to have a Hill-type dependence on the cytosol Ca^2+^ level^106^, and is given by F_*S*_ = V_*S*_[Ca^2+^]^2^/([Ca^2+^]^2+^ K_*S*_^2^), with the half-activation constant K_*S*_ set to 0.2 μM^113^. As with the plasma membrane efflux pumps, we introduce a compensatory leak term at the ER surface, k_*S*_([Ca^2+^]_*ER*_-[Ca^2+^]), that balances the SERCA pump activity to help maintain a resting cytosolic [Ca^2+^] = 50 nM under basal conditions. The magnitude of maximal SERCA activity in the spine, V_*S*_, depends on the density of SERCA molecules on the ER membrane, ER surface area in the spine head, and pump efficiency per molecule. As no estimate appropriate for a spine-sized region was readily available, we have obtained an approximate estimate for V_*S*_ by referring to previous experimental measurements on ER refilling rates in rat sensory neuron^111, 114^. The refilling time constant there was found to be about 1-5 min. By making reasonable assumptions about the total cell volume, ER volume fraction, SERCA surface density (β2000 μm^21-15^), and spine ER area (assumed to be 10% of the spine head area^29^), we have arrived at an estimate of V_*S*_ =1 μM/s, which sets the scale for the contribution of SERCA activity to Ca^2+^ sequestration in a typical ER+ spine head.

### Modeling synaptic activation

Unitary synaptic input at the spine head is modeled as a single pulse of glutamate, which transiently binds to postsynaptic mGluR, and concurrently activates AMPAR/NMDAR currents as described above to produce a small (~3-5 mV) depolarization of the postsynaptic membrane (EPSP). The amplitude of spine Ca^2+^ evoked by NMDAR current during an EPSP (Δ*Ca_EPSP_*) is considered to be in the range of a few hundred nM. For our control synapse (ER-spine) we have set the NMDAR conductance parameter g_*N*_ to 65 pS, which yields a Δ*Ca_EPSP_* = 0.2 μM, consistent with previous studie^51, 52^. In order to assess the parameter dependence of our results, we have also varied g_*N*_ to obtain a range of Δ*Ca_EPSP_* values (0.1-1 μM). The delay in Ca^2+^ release from the ER following the NMDAR-mediated Ca^2+^ rise is measured with respect to the time of application of glutamate.

Induction of frequency-dependent plasticity in ER-/ER+ spines is simulated with trains of regularly spaced synaptic inputs applied over a range of different frequencies (0.1-20 Hz). In order to simulate the activation of Schaffer collateral (SC) fibers during the induction of LTP/LTD, we assume that the local dendritic shaft (modeled as a passive compartment) receives synchronous input from multiple synapse^67, 116^, which amplifies the depolarization at the spine membrane through the passive resistive coupling (*R_C_*), leading to stronger activation of the NMDAR current at the spine head. The parameter *ρ_S_*, which sets the magnitude of the total number of co-active synaptic inputs onto the dendritic compartment during SC stimulation, has been adjusted to evoke a depolarization of ~10 mV at the spine head in response to a single input^49^. As the modest depolarization of the spine in this setting is insufficient for L-VGCC activation, their contribution to the Ca^2+^ response is ignored for simplicity.

We also model the pairing of synaptic inputs with the strong depolarization of the postsynaptic membrane induced by backpropagating action potentials (BAPs) in the CA1 neuron, mimicking the conditions for the induction of spike timing-dependent plasticity (STDP)^60^, which involves the activation of both the NMDAR and L-VGCC. Dendritic BAPs are modeled as a voltage profile with a peak depolarization of Vo = 67 mV, and composed of a fast (*τ_f_* = 3 ms) and a slow (*τ_s_* = 40 ms) exponentially decaying component^51, 52^:

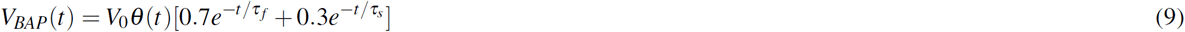

where *θ*(t) = 1 for t ≥ 0, and 0 otherwise, for a BAP arriving at t=0. A BAP is simulated by feeding the above voltage profile to the dendritic compartment. During repetitive stimulation, each synaptic input is paired with either 1 or 2 BAPs (spike doublets/triplets). The timing difference between the pre and postsynaptic firing, Δt, is measured as the interval between the start of the glutamate pulse and peak of the BAP at the dendritic compartment. When a synaptic input is paired with a postsynaptic burst composed of two BAPs (spike triplet), Δt is defined as the interval between the glutamate pulse and the peak of the *second* BAP; BAPs in a burst are separated by a fixed interval of 10 ms. By convention, glutamate release preceding the BAP is assigned a positive Δ*t*.

### Ca^2+^**-based plasticity model**

Induction of long-term modification at hippocampal/cortical synapses is governed by the magnitude and duration of local Ca^2+^ elevation at the dendritic spine^117^. Long-term potentiation (LTP) is found to be reliably induced with short volleys of high-frequency stimulation^118^, whereas synaptic depression (LTD) usually requires sustained stimulation at low rates lasting several minutes^56^. We adopt a previously proposed model^49^ for bidirectional plasticity to describe the induction of both rate and spike timing-dependent synaptic efficacy changes at the ER-bearing spine (Fig. 1D). The model describes the dynamics of a (dimensionless) weight variable, *w* (a proxy for the postsynaptic AMPAR conductance), and its functional form approximates biophysically detailed descriptions of the regulation of AMPAR number and/or phosphorylation level by a combination of Ca^2+^-activated kinases and phosphatase^119, 120^. Following some earlier studie^67, 97, 98^, we model the dependence of w on the concentration of active calmodulin (aCaM) instead of the free Ca^2+^. Ca^2+^-bound CaM is known to regulate the activation of several downstream effectors such as Ca^2+^/CaM-dependent protein kinase II (CaMKII)^121^, calcineurin^122^, and protein kinase^123, 124^, which converge to mediate synaptic changes underlying early LTP/LTD expressio^125-128^. Although earlier models assumed that only the fully activated form of CaM (bound to 4 Ca^2+^ molecules) is relevant, recent studies indicate that even the partially bound forms are capable of allosteric activity (e.g., activation of CaMKII^121^); thus, as a measure of CaM activity we consider the total Ca^2+^-bound CaM in the spine. The dynamic range of aCaM is restricted by the availability of total CaM in the spine, unlike the Ca^2+^ response which is not bounded and in principle can grow very large during persistent high frequency stimulation. The dynamical equation governing the Ca^2+^ dependence of *w* is given by

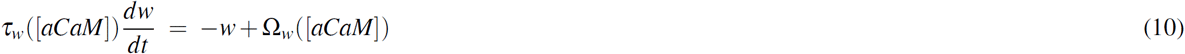

where the function Ω_*w*_ is modeled as the difference between two sigmoid functions^49^:

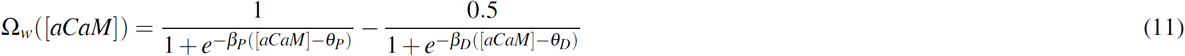

with slopes *β_D_* = 60 and *β_P_* = 60. The offsets *θ_P_* and *θ_D_* (*θ_P_* > *0_D_*) control the thresholds for the induction of LTP/LTD during synaptic stimulation: no plasticity is induced when aCaM levels remain below *θ_D_*, LTD is induced when aCaM is restricted by and large to the interval (*θ_D_,θ_P_*), and LTP induction requires higher levels of aCaM, exceeding the threshold *θ_P_*. The temporal factor *τ_w_* is given by

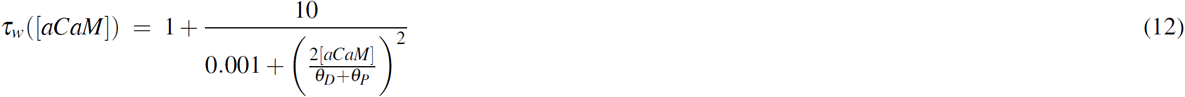

and its parameters have been set in accordance with experimentally suggested rates for the induction of early LTP (βseconds) and LTD (~ minutes) and the persistence of synaptic weight changes under resting conditions (~1-3 h)^49^.

Rate-dependent plasticity is simulated over the 0.1-20 Hz frequency range (in steps of 0.1 Hz) with 900 presynaptic SC inputs applied at each frequency, following earlier studie^13, 49, 56^. STDP induction is mimicked with a train of 100 pairings of pre/postsynaptic spiking presented at 5 Hz^60^, with the spike timing difference Δ*t* varied from-100 ms and 100 ms in steps of 1 ms. The weight variable *w* (assumed to be 0 initially in all our simulations) integrates over the temporal spine Ca^2+^ signal evoked by these synaptic activation patterns, resulting in a net (cumulative) change Δw at the end of the stimulation period. Spine ER is assumed to contribute to the Ca^2+^ pool in the spine driving changes in *w*, thereby modulating the induction of NMDAR-dependent plasticity and giving rise to possible differences in Δ*w* relative to the reference ER-spine. We characterize the nature of this differential effect of spine ER, ΔΔ*w*, as a function of various biophysical model parameters. The plasticity profile obtained in our model synapse is shaped by the choice of *θ_P_/θ_D_* (assumed to be same for the ER+ and ER-spines), which are appropriately set to have agreement with experimentally obtained plasticity curves. For the rate-dependent plasticity, *θ_P_* is adjusted such that LTP induction occurs for frequencies ≥ 15 Hz in the ER-spine head^13, 56^(although it represents an average obtained from a large number of synapses, we assign this threshold to every individual synapse); we also examine the robustness of our results to variation in this parameter (*f_P_* = 10-20 Hz). Different choices of the LTD threshold (*θ_D_*) for the ER-spine have been similarly explored (*f_D_* = 1-6 Hz). In the case of STDP, we refer to a previous experimental stud^60^ which reports the average plasticity curve obtained from measurements at a population of CA3-CA1 synapses; here, again, we ascribe the plasticity thresholds (Δ*t_D_*, Δ*t_P_*) read off from the average curve to our reference synapse associated with an ER-spine (implicitly assuming that only a minor proportion of the synapses recorded from are associated with an ER). We also follow this up by examining the sensitivity of the results to variation in Δ*t_D_* and Δ*t_P_* over ±10 ms windows.

All numerical simulations and analysis comprising this study were carried out using Python.

## Acknowledgements

Financial support from Wellcome Trust/DBT India Alliance (S.N.) and Science & Engineering Research Board, India (G.M.) is gratefully acknowledged.

## Data/code availability

All information required to replicate the presented results is included in the article and supplementary material. Python code for running the simulations is available from the authors on request.

## Author contributions statement

G.M.: Conceptualization, Methodology, Investigation, Formal analysis, Visualization, Writing-Original Draft Preparation, Writing-Review and Editing.

S.N.: Conceptualization, Methodology, Investigation, Writing-Review and Editing, Funding Acquisition.

## Supporting Information (SI)

### Effect of varying SERCA pump activity on Ca^2+^ handling in ER+ spines

Besides providing an additional source of Ca^2+^ in the spine head, ER also contributes to spine Ca^2+^ regulation via the SERCA pumps that extrude Ca^2+^ from the cytosol and are responsible for maintaining the steep (β1000-fold) Ca^2+^ concentration gradient across the ER membrane. In all our simulations, the SERCA activity in the ER+ spine was set to a fixed value (V_*S*_ = 1*μ*M/s), which was inferred (under biologically realistic assumptions) from whole-cell experimental data on ER refilling rates (Methods). This estimate of SERCA activity has only a minor effect on the decay rate of spine Ca^2+^ signals, which is primarily determined by the efficiency of Ca^2+^ transport across the plasma membrane (mediated by PMCA and NCX). However, some previous studies suggest a more prominent role for SERCA in regulating the time course of Ca^2+^ responses and its clearance from the spine head (Sabatini et al., 2002; Bell et al., 2018). Given this context, we asked how higher levels of SERCA activity would modulate the Ca^2+^ signals evoked by synaptic activation in an ER+ spine head.

We simulated the responses of our synaptic model to a range of synaptic input frequencies (0.1-20 Hz), with varying levels of SERCA activity parametrized by *V_S_*. Fig. S6A compares the frequency-response relation of cytosolic Ca^2+^-CaM in the ER-spine (black) with an equivalent ER+ spine with 30IP_3_R and *V_S_* spanning 3 orders of magnitude (colored curves). In general, ICCR significantly augments the NMDAR-driven Ca^2+^ elevation at low frequencies; however, the magnitude and frequency window of enhancement of Ca^2+^ responses is attenuated at higher SERCA efflux rates. ICCR is progressively suppressed with increasing frequency due to the Ca^2+^-dependent inhibition of IP_3_R, and at high enough frequencies, SERCA pump activity could potentially counterbalance the contribution of ICCR, resulting in a net *reduction* in the total Ca^2+^/CaM response in the ER+ spine, relative to the ER-spine. This becomes more apparent at higher VS, and is illustrated in Fig. S6A by the Ca^2+^/CaM profiles for V_S_ = 100-1000 *μ*M/s. Reflecting these trends, the corresponding plasticity curves in Fig. S6B suggest a complex, frequency-dependent modulation of plasticity, the exact nature of which is determined by an interplay between number of IP_3_R, SERCA efflux rate, and the NMDAR activation profile. A direct consequence of reduced Ca^2+^ elevation in the ER+ spine (relative to the ER-spine) at high SERCA efflux rates is a shift in the threshold for LTP induction to higher input frequencies. At moderate *V_S_* values (e.g., 100 *μ*M/s), enhanced LTD at low frequencies is accompanied by extension of the LTD window and rightward shift of *f_P_*. At higher *V_S_* values (≳ 500 *μ*M/s), however, LTD can be suppressed as well, resulting in an overall shift of the entire plasticity curve to higher induction frequencies.

In summary, our results indicate that the overall contribution of ER to spine Ca^2+^ regulation is shaped by an interplay between Ca^2+^ release (ICCR) and buffering/uptake (SERCA), and this becomes particularly relevant at high SERCA activity levels. Clarity on the nature and extent of modulation of spine Ca^2+^ signaling by Ca^2+^ uptake into ER will require more precise biochemical quantification of SERCA kinetics, and its subcellular distribution in pyramidal cell dendrites.

---

**Table S1.** Parameters governing membrane voltage dynamics.

| Parameter | Symbol | Value |
| --- | --- | --- |
| Avogadro number | $N_a$ | $6.023 \times 10^{23}/\text{mole}$ |
| Electric charge per $\text{Ca}^{2+}$ ion | $q_{Ca}$ | $3.2 \times 10^{-19} \text{ C}$ |
| Volume of (spherical) ER-/ER+ spine head | $V_{spine}$ | $0.06 \mu\text{m}^3$ |
| Resting membrane potential in spine/dendrite | $u_{rest}$ | -70 mV |
| Resistive coupling between spine head and parent dendrite | $R_C$ | 100 MΩ |
| AMPA conductance parameter | $g_A$ | 0.5 nS |
| AMPA rise time constant | $\tau_A^r$ | 0.2 ms |
| AMPA decay time constant | $\tau_A^d$ | 2 ms |
| AMPA current reversal potential | $E_A$ | 0 mV |
| NMDAR conductance parameter | $g_N$ | 65 pS in model synapse (gives $\Delta Ca_{EPSP} = 0.2 \mu\text{M}$) |
| NMDAR rise time constant | $\tau_N^r$ | 5 ms |
| NMDAR decay time constant | $\tau_N^d$ | 50 ms |
| NMDAR current reversal potential | $E_N$ | 0 mV |
| VGCC activation offset | $u_m$ | -20 mV |
| VGCC activation slope | $k_m$ | 5 mV |
| VGCC activation time constant | $\tau_m$ | 0.08 ms |
| VGCC inactivation offset | $u_h$ | -65 mV |
| VGCC inactivation slope | $k_h$ | -7 mV |
| VGCC inactivation time constant | $\tau_h$ | 300 ms |
| Maximum depolarization during BAP in dendritic compartment | $V_0$ | +67 mV |
| Fast time constant of BAP decay | $\tau_f$ | 3 ms |
| Slow time constant of BAP decay | $\tau_s$ | 40 ms |
| Effective number of co-active synaptic inputs at dendritic compartment (rate-dependent plasticity) | $\rho_S$ | $5 \times 10^5 \text{ cm}^{-2}$ |

**Table S2.** Reaction rate parameters for Ca^2+^ buffering and extrusion.

| Parameter | Symbol | Value |
| --- | --- | --- |
| Kinetic rate constants for interaction between $\text{Ca}^{2+}$ and calbindin (CB) | $k_{M0M1}$ | $174 \mu\text{M}^{-1}\text{s}^{-1}$ |
| | $k_{M1M2}$ | $87 \mu\text{M}^{-1}\text{s}^{-1}$ |
| | $k_{M1M0}$ | $35.8 \text{s}^{-1}$ |
| | $k_{M2M1}$ | $71.6 \text{s}^{-1}$ |
| | $k_{H0H1}$ | $22 \mu\text{M}^{-1}\text{s}^{-1}$ |
| | $k_{H1H2}$ | $11 \mu\text{M}^{-1}\text{s}^{-1}$ |
| | $k_{H1H0}$ | $2.6 \text{s}^{-1}$ |
| | $k_{H2H1}$ | $5.2 \text{s}^{-1}$ |
| Kinetic rate constants for interaction between $\text{Ca}^{2+}$ and endogenous immobile buffer (CBP) | $k_f^{CBP}$ | $247 \mu\text{M}^{-1}\text{s}^{-1}$ |
| | $k_b^{CBP}$ | $524 \text{s}^{-1}$ |
| Kinetic rate constants for interaction between $\text{Ca}^{2+}$ and endogenous slow buffer | $k_f^{slow}$ | $24.7 \mu\text{M}^{-1}\text{s}^{-1}$ |
| | $k_b^{slow}$ | $52.4 \text{s}^{-1}$ |
| Kinetic rate constants for interaction between $\text{Ca}^{2+}$ and calmodulin (CaM) | $k_{C0C1}$ | $6.8 \mu\text{M}^{-1}\text{s}^{-1}$ |
| | $k_{C1C2}$ | $6.8 \mu\text{M}^{-1}\text{s}^{-1}$ |
| | $k_{C1C0}$ | $68 \text{s}^{-1}$ |
| | $k_{C2C1}$ | $10 \text{s}^{-1}$ |
| | $k_{N0N1}$ | $108 \mu\text{M}^{-1}\text{s}^{-1}$ |
| | $k_{N1N2}$ | $108 \mu\text{M}^{-1}\text{s}^{-1}$ |
| | $k_{N1N0}$ | $4150 \text{s}^{-1}$ |
| | $k_{N2N1}$ | $800 \text{s}^{-1}$ |
| Kinetic rate constants for PMCA pump | $k_f^{PMCA}$ | $150 \mu\text{M}^{-1}\text{s}^{-1}$ |
| | $k_b^{PMCA}$ | $15 \text{s}^{-1}$ |
| | $k_3^{PMCA}$ | $12 \text{s}^{-1}$ |
| | $k_L^{PMCA}$ | $3.33 \text{s}^{-1}$ |
| Kinetic rate constants for NCX | $k_f^{NCX}$ | $300 \mu\text{M}^{-1}\text{s}^{-1}$ |
| | $k_b^{NCX}$ | $300 \text{s}^{-1}$ |
| | $k_3^{NCX}$ | $600 \text{s}^{-1}$ |
| | $k_L^{NCX}$ | $10 \text{s}^{-1}$ |

**Table S3.**
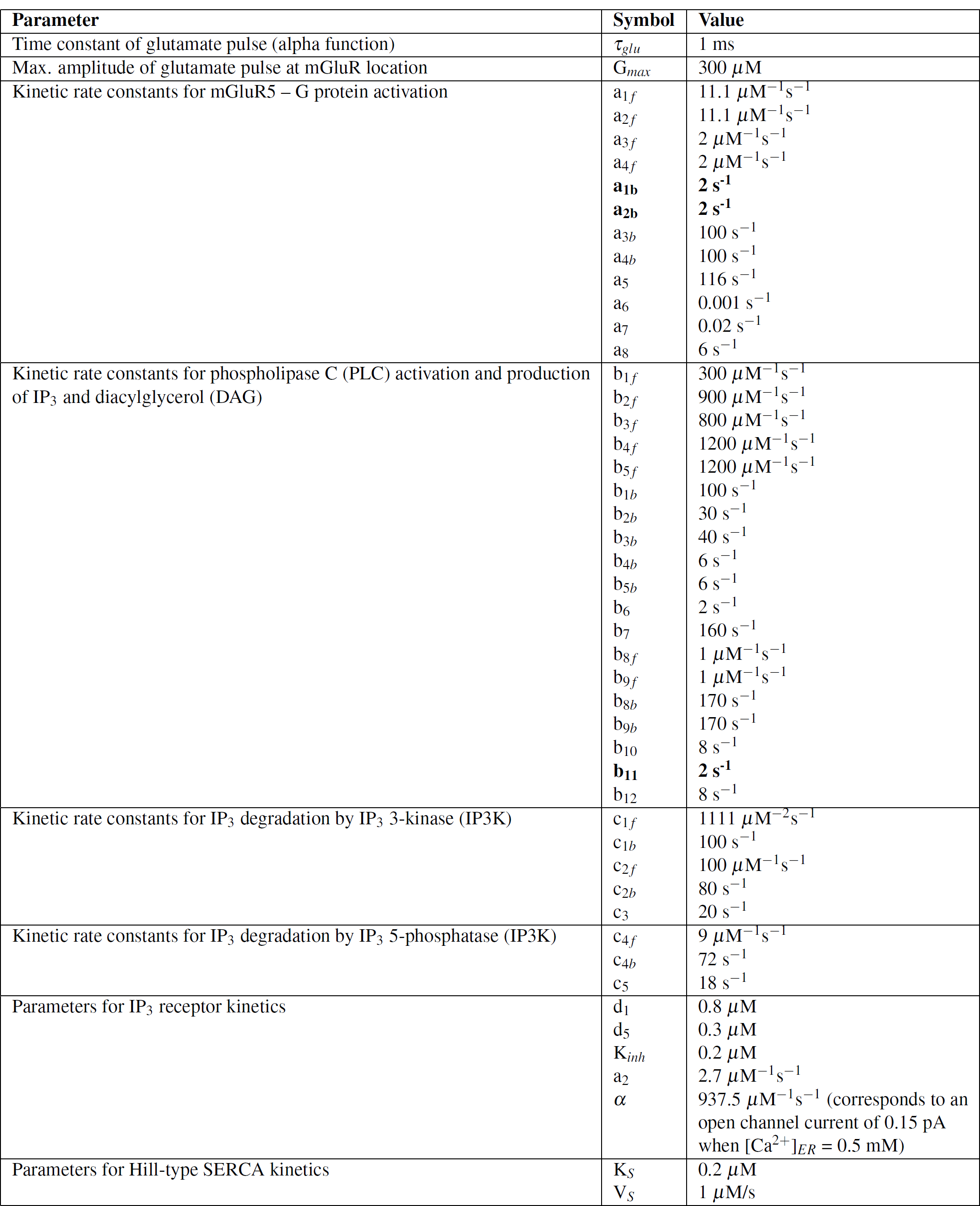
Reaction parameters for mGluR-IP3 signaling and ER Ca^2+^ handling. (Parameter values which have been changed from the original reference^54^ are highlighted.)

**Table S4.** Concentrations of various chemical species in the model.

| Species | Value |
| --- | --- |
| Resting $\text{Ca}^{2+}$ concentration in the spine cytosol ($[\text{Ca}^{2+}]_{rest}$) | 50 nM |
| Resting $\text{IP}_3$ concentration in the spine | 0.1 $\mu\text{M}$ |
| $\text{Ca}^{2+}$ concentration in ER lumen (fixed) | 250 $\mu\text{M}$ |
| Fixed extracellular $\text{Ca}^{2+}$ concentration ($[\text{Ca}^{2+}]_{ext}$) | 2 mM |
| Total calbindin (CB) concentration | 45 $\mu\text{M}$ |
| Total concentration of endogenous immobile buffer (CBP) | 80 $\mu\text{M}$ |
| Total concentration of endogenous slow buffer | 40 $\mu\text{M}$ |
| Total calmodulin (CaM) concentration | 50 $\mu\text{M}$ |
| Surface density of PMCA pumps | 1000 $\mu\text{m}^{-2}$ |
| Surface density of NCX pumps | 140 $\mu\text{m}^{-2}$ |
| Total mGluR concentration in the spine | 0.3 $\mu\text{M}$ |
| Total PLC-$\text{PIP}_2$ concentration in the spine | 0.8 $\mu\text{M}$ |
| Total $\text{PIP}_2$ concentration | 4 mM |
| Total Gq-GDP concentration in the spine | 1 $\mu\text{M}$ |
| Total $\text{IP}_3$ 3-kinase (IP3K) in the spine | 0.9 $\mu\text{M}$ |
| Total $\text{IP}_3$ 5-phosphatase (IP5P) in the spine | 1 $\mu\text{M}$ |
| Number of $\text{IP}_3$ receptors associated with spine ER ($N_R$) | 10-50 |

**Figure S1.**
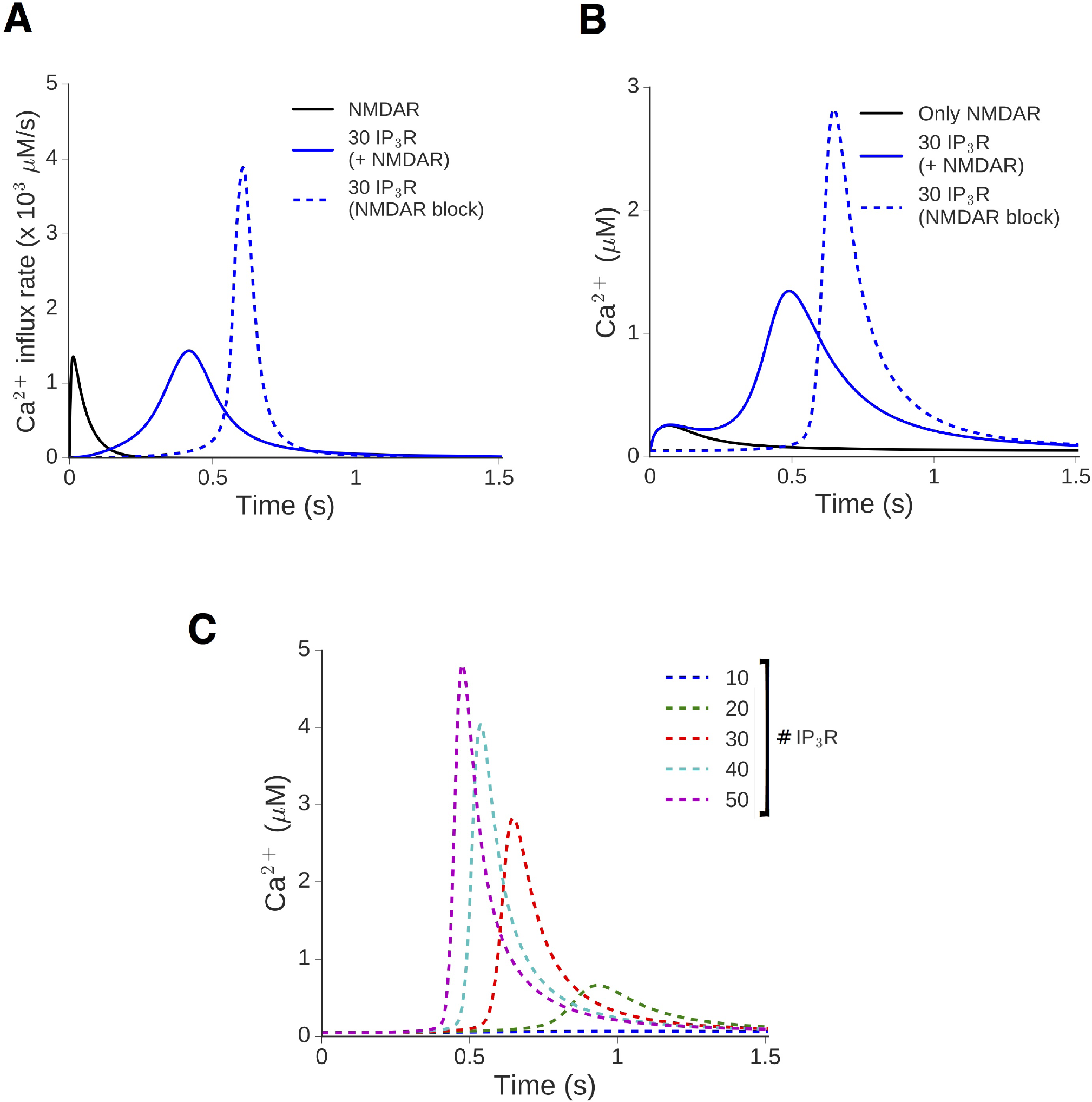
Store Ca^2+^ release evoked by mGluR signaling does not require the initial NMDAR-mediated Ca^2+^. Store Ca^2+^ release evoked by mGluR signaling does not require the initial NMDAR-mediated Ca^2+^ transient. Illustrative responses in our model of a CA1 spine head to a pulse of glutamate applied at t=0 (Δ*Ca_EPSP_* = 0.2 μM). **A**, Time course of the Ca^2+^ influx rate through NMDAR-gated channels (black), and IP_3_R-gated channels in the presence (blue, solid curve) and absence (blue, dashed curve) of the NMDAR. **B**, The Ca^2+^ traces in the ER-spine with only NMDAR (black), in the ER+ spine with both NMDAR and IP3R present (blue, solid curve), and in the ER+ spine with NMDAR blocked (blue, dashed curve). **C**, Ca^2+^ transients in an ER+ spine with NMDAR blocked, for different numbers of IP3R (10-50).

**Figure S2.**
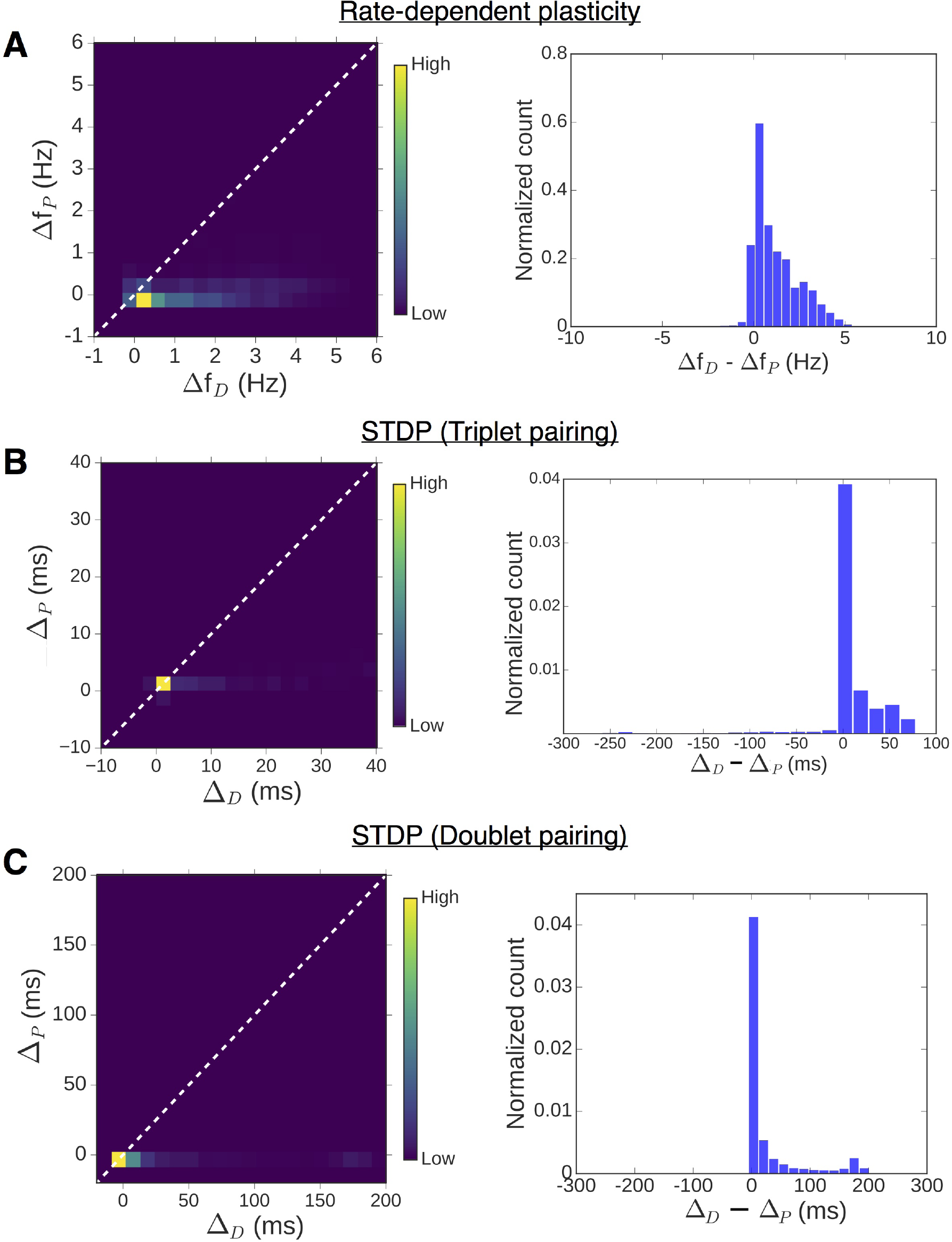
Differential enhancement of LTD and LTP by spine ER is broadly reproduced over a range of realistic parameter settings in our model. Summary statistics of changes in plasticity windows introduced by ICCR in the case of frequency-dependent plasticity (A), and for STDP with triplet (B) and doublet (C) spike pairing stimuli. In each case, the left figure is the 2D heat map of the distribution obtained from random sampling of the thresholds at the ER-control spine, over a range of IP_3_R cluster sizes (10-50); the same information is represented as a 1D histogram of the relative differences (depression v/s potentiation) in the right figure. (For frequency-dependent plasticity, *f_D_* and *f_p_* have been randomly sampled from 1-6 Hz and 10-20 Hz ranges, respectively. For STDP, Δ*t_D_* and Δ*t_P_* have been sampled from 20 ms windows centered on-35 ms and +35 ms, respectively.)

**Figure S3.**
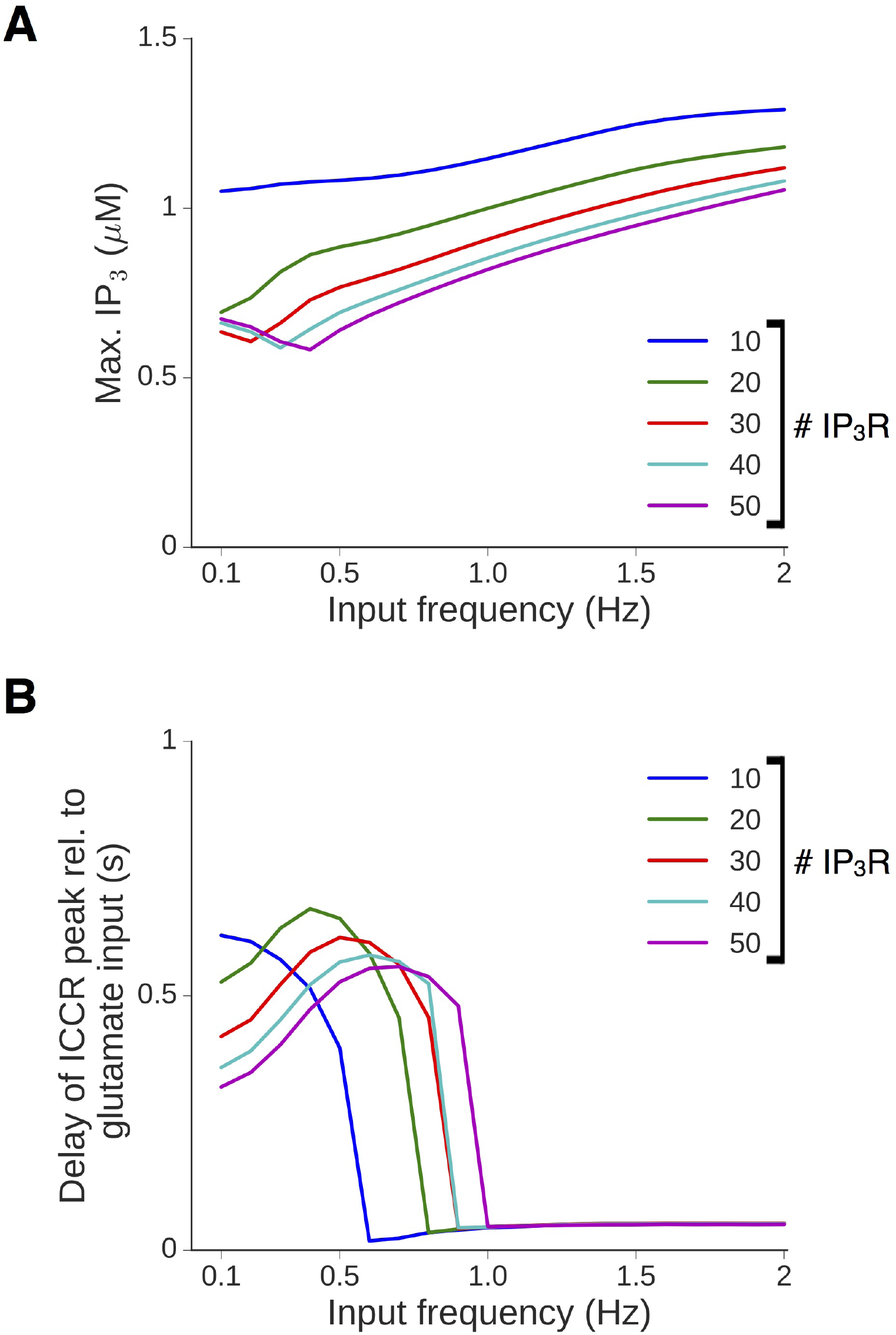
Dependence of ICCR time course on the frequency of input at low frequencies (0.1-2 Hz). **A**, Maximum IP3 level attained during persistent synaptic stimulation as a function of the stimulus frequency, for different IP_3_R cluster sizes. **B**, Dependence of the lag in peak IP_3_R Ca^2+^ current (relative to the glutamate input) on the stimulus frequency, for different IP_3_R cluster sizes. (All results for a spine head with Δ*Ca_EPSP_* = 0.2 μM.)

**Figure S4.**
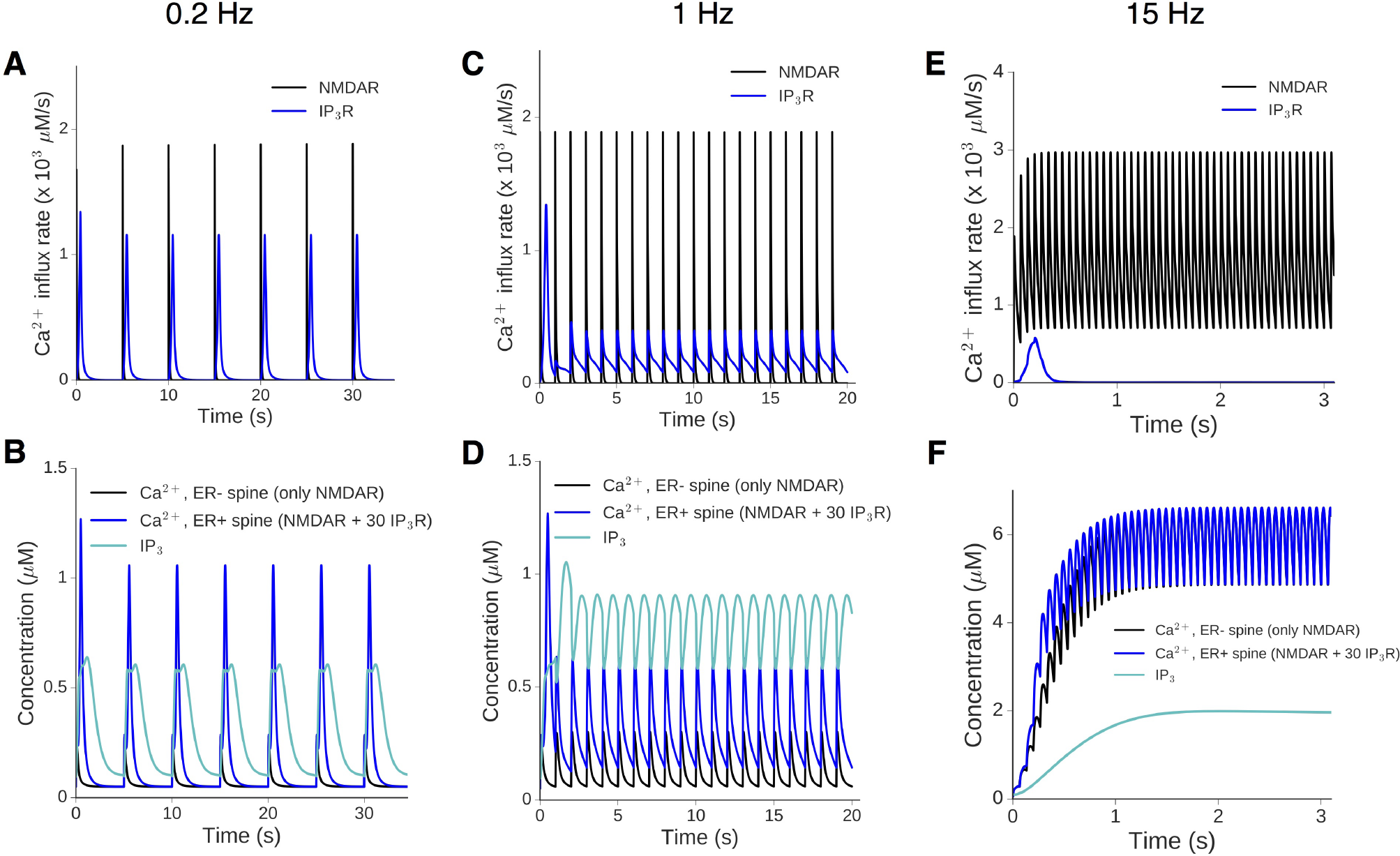
Dependence of store Ca^2+^ release on the frequency of synaptic input. Representative traces of the Ca^2+^ response in the ER-and ER+ spines at three different input frequencies: f = 0.2 Hz (A,B), 1 Hz (C,D) and 15 Hz (E,F). In each case, the top figure shows the Ca^2+^ current through the NMDAR-gated (black) and IP_3_R-gated channels (blue), while the bottom figure displays the Ca^2+^ trace in the ER-(black) and ER+ (blue) spines along with the time course of IP_3_ in the spine head (cyan). At low frequencies (0.2 and 1 Hz), every input evokes release from stores which is comparable to the NMDAR-mediated Ca^2+^ transient. At higher frequencies (15 Hz), sustained NMDAR-mediated Ca^2+^ elevation inhibits the IP_3_R (E) and there is near-convergence of the steady-state Ca^2+^ time courses in the ER+ and ER-spines.

**Figure S5.**
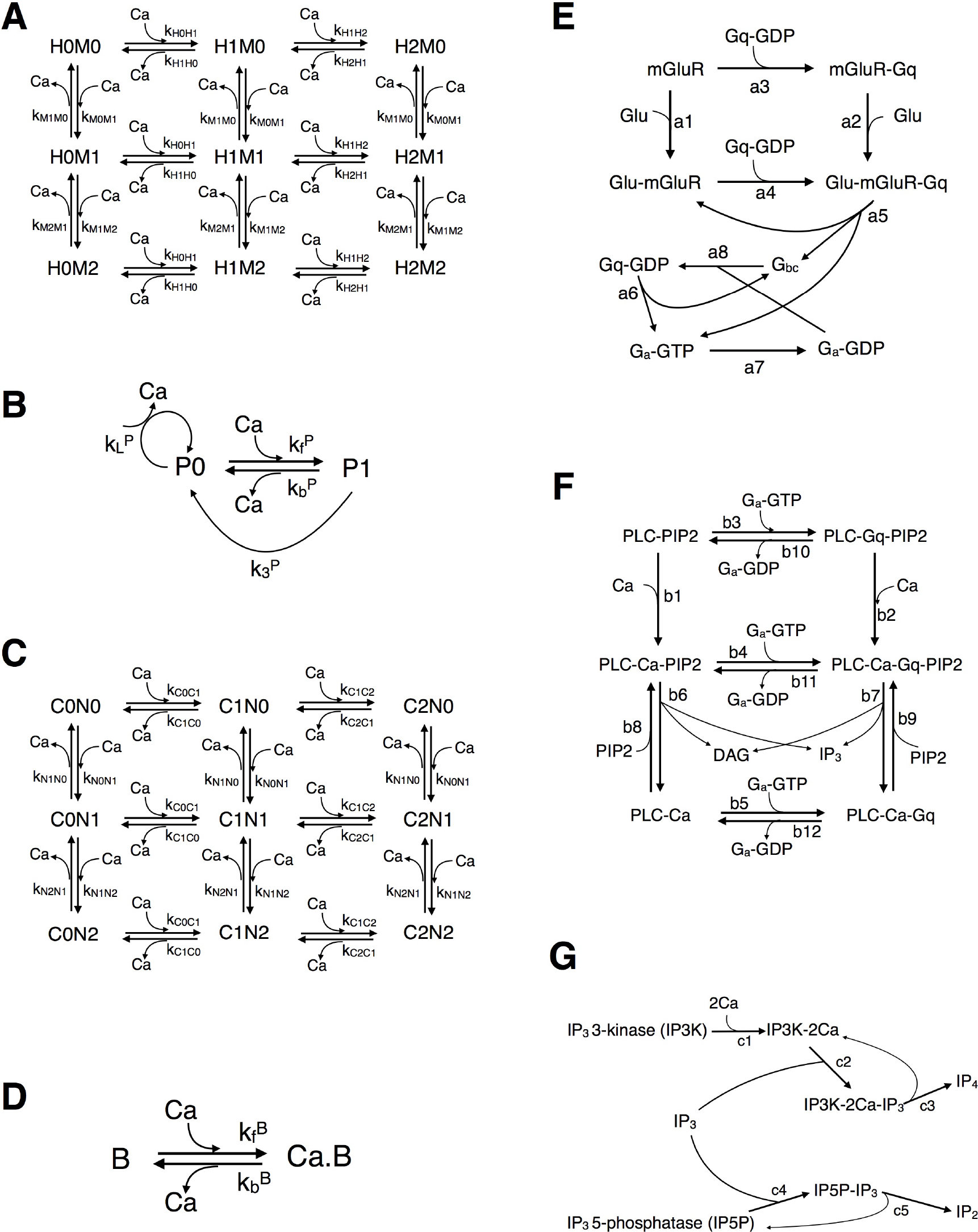
Reaction kinetics for various chemical species in the model. **A**, Nine-state kinetic scheme for Ca^2+^ interaction with calbindin (CB). Each CB molecule possesses two medium affinity (M) and two high affinity (H) Ca^2+^ binding sites. **B**, First-order reversible reaction describing the action of endogenous immobile and slow Ca^2+^ buffers. **C**, Reaction network for the interaction of Ca^2+^ with the N and C lobes of calmodulin (CaM). **D**, Michaelis-Menten kinetic scheme to describe the action of plasma membrane Ca^2+^ pumps (P = PMCA or NCX). E, Biochemical model for G-protein (Gq) activation by group I metabotropic glutamate receptor (mGluR). **F**, Detailed model describing the G-protein-mediated activation of PLC and subsequent production of IP_3_ and DAG via hydrolysis of PIP_2_. G, Reaction steps involved in the enzymatic degradation of IP_3_ by IP_3_ 3-kinase (IP3K), which has Ca^2+^-dependent activity, and IP_3_ 5-phosphatase (IP5P). (Note: Several reaction steps in **E-G** are bidirectional (reversible), but have been represented as arrows in a single direction to avoid clutter. The full set of forward and backward kinetic rate constants is listed in Tables S2 and S3.)

**Figure S6.**
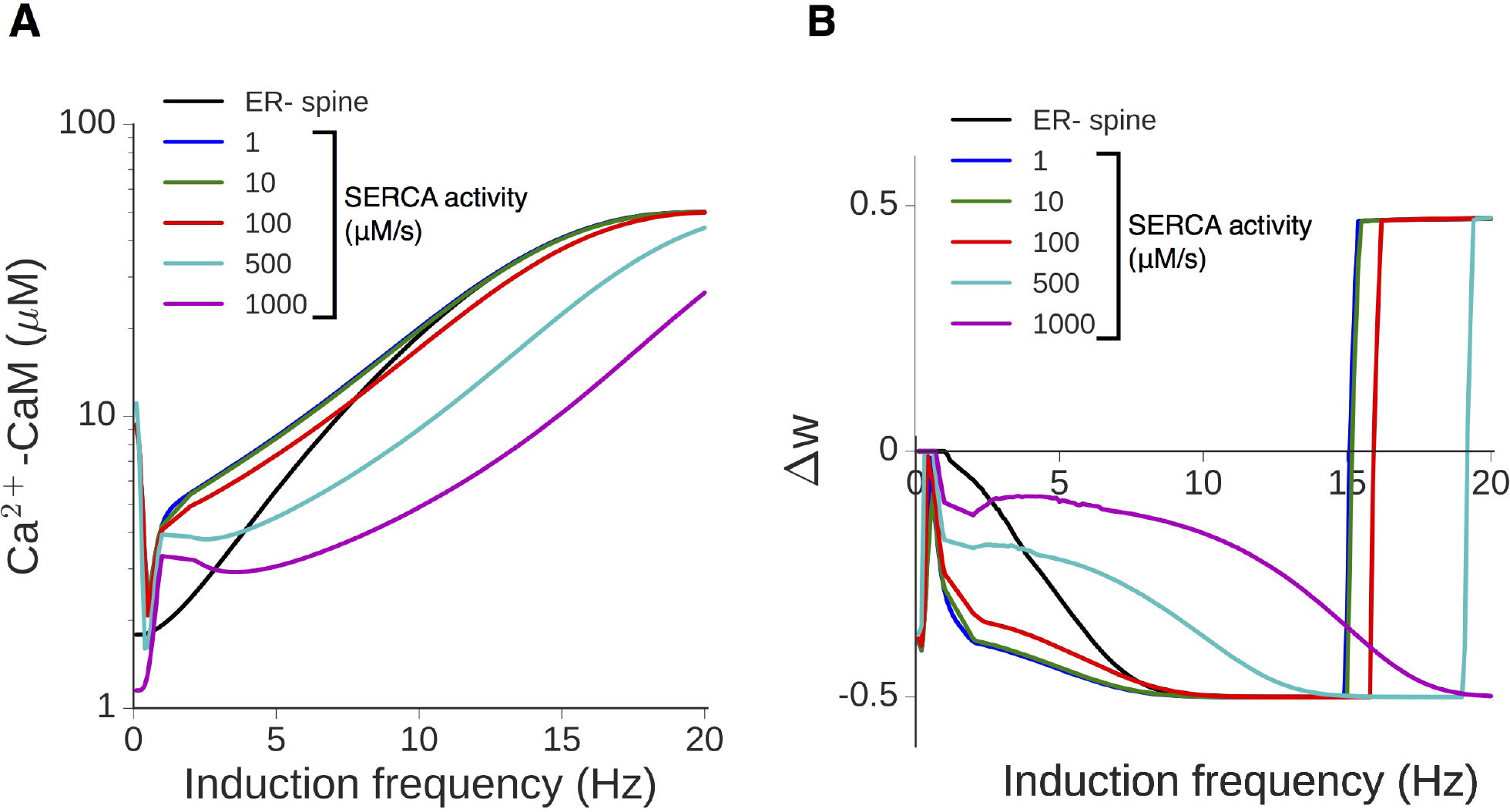
Increased SERCA activity in the spine can modulate the threshold for LTP induction. **A**, Dependence of CaM activation on the synaptic input frequency for the ER-control spine (black) and ER+ spines with increasing levels of SERCA extrusion rate (colored curves); N_*R*_ is set to 30 here, and Δ*Ca_EPSP_* = 0.2 μM in the ER-spine. **B**, The corresponding plasticity profiles (total weight change at the end of 900 input spikes).

